# The dynamics and longevity of circulating CD4^+^ memory T cells depend on cell age and not the chronological age of the host

**DOI:** 10.1101/2023.10.16.562650

**Authors:** M. Elise Bullock, Thea Hogan, Cayman Williams, Sinead Morris, Maria Nowicka, Minahil Sharjeel, Christiaan van Dorp, Andrew J. Yates, Benedict Seddon

**Author notes:** These authors contributed equally to this work (M. Elise Bullock, Thea Hogan, Cayman Williams). These authors also contributed equally to this work (Andrew J. Yates, Benedict Seddon). Address correspondence to either author.

## Abstract

Quantifying the kinetics with which memory T cell populations are generated and maintained is essential for identifying the determinants of the duration of immunity. The quality and persistence of circulating CD4^+^ effector memory (T_EM_) and central memory (T_CM_) T cells in mice appear to shift with age, but it is unclear whether these changes are driven by the aging host environment, by cell age effects, or both. Here we address these issues by combining DNA labelling methods, established fate-mapping systems, a novel reporter mouse strain, and mathematical models. Together, these allow us to quantify the dynamics of both young and established circulating memory CD4^+^ T cell subsets, within both young and old mice. We show that that these cells and their descendents become more persistent the longer they reside within the T_CM_ and T_EM_ pools. This behaviour may limit memory CD4 T cell diversity by skewing TCR repertoires towards clones generated early in life, but may also compensate for functional defects in new memory cells generated in old age.

**Author summary:** Our long-term protection against infections depends in part on the maintenance of diverse populations of memory CD4 T cells, which are made in response to the initial exposure to the pathogen or a vaccine. These cells are not long-lived, but instead are maintained dynamically at a clonal level through loss and division. Understanding how immune memory persists therefore requires measuring these rates of these processes, and how they might change with age. Here we combine experiments in mice with mathematical models to show that memory CD4 T cells exhibit complex dynamics but increase their capacity to survive as they age. This dynamic implies that as individuals age, their memory CD4 T cell populations become enriched for older clones. This established memory may compensate for functional defects in new T cell responses generated later in life.

## Introduction

CD4 T cells can suppress pathogen growth directly and coordinate the activity of other immune cells [1, 2]. Following exposure to antigen, heterogeneous populations of circulating memory CD4 T cells are established that enhance protection to subsequent exposures. In mice, these can be categorised broadly as central (T_CM_) or effector (T_EM_) memory. Canonically defined T_CM_ have high expression of CD62L, which allow them to circulate between blood and secondary lymphoid organs, while CD4 T_EM_ express low levels of CD62L and migrate to tissues and sites of inflammation. The lineage relationships between CD4 T_CM_, T_EM_, and other memory subsets remain unclear [3].

In mice, circulating memory CD4 T cells appear not to be intrinsically long-lived on average, but are lost through death or onward differentiation on timescales of days to weeks [4–9]. However, compensatory self-renewal can act to sustain memory T cell clones, defined as populations sharing a given T cell receptor (TCR), for much longer than the lifespan of their constituent cells [10]. To understand in detail how the size of memory T cell populations change with time since exposure to a pathogen, we therefore need to be able to quantify their rates of *de novo* production, division, and loss, any heterogeneity in the rates of these processes, how kinetically distinct subsets are related, and how their kinetics might change with host and/or cell age.

Much of our knowledge of memory CD4 T cell dynamics derives from studies of laboratory mice housed in specific pathogen-free conditions who have not experienced overt infections, but nevertheless have abundant memory phenotype (MP) CD4 T cells [11–13]. These populations contain both rapidly dividing and more quiescent populations [4, 14–17], a stratification that holds within the T_CM_ and T_EM_subsets [5]. MP T cells are established soon after birth, and attain broadly stable numbers determined in part by the cleanliness of their housing conditions [18]. After the first few weeks of life, however, both CD4 T_CM_ and CD4 T_EM_ are continuously replaced at a rates of a few percent of the population size per day [5], independent of their housing conditions [18]. MP CD4 T cells are therefore likely specific for an array of commensals, self, or ubiquitous environmental antigens, and may comprise subpopulations at different stages of maturation or developmental end-points. Dissecting their dynamics in detail may then shed light on the life-histories of conventional memory cells induced by infection.

A widely-used approach to studying lymphocyte population dynamics is DNA labelling, in which an identifiable agent such as bromodeoxyuridine (BrdU) or deuterium is taken up during cell division. By measuring the frequency of cells within a population that contain the label during and following its administration, one can use mathematical models to extract rates of production and turnover (loss). Without allowing for the possibility variation in these rates within a population, their average estimates may be unreliable [19, 20]. Explicit treatment of such ‘kinetic heterogeneity’ can yield estimates of average production and loss rates that are interpretable and independent of the labelling period, although the number of distinct subsets and the lineage relationships between them are difficult to resolve [4, 5, 18, 21, 22]. Further, models of labelling dynamics usually conflate the processes of influx of new cells and self-renewal of existing ones into a single rate of cell production. Distinguishing these processes is particularly important in the context of memory, because their relative contributions determine the persistence of clones within a population; for a population at equilibrium, clones are diluted by influx, but sustained by self-renewal.

To address these uncertainties, one can combine DNA labelling with information from other systems to validate predictions or constrain the choice of models. One approach is to obtain an independent estimate of any constitutive rate of flow into a population by following its replacement by labelled precursors. In a series of studies, we established a mouse model that yields the rates of replacement of all peripheral lymphocyte subsets at steady state [5, 18, 23–28]. In this system, illustrated in Fig 1A, low doses of the transplant conditioning drug busulfan are administered to specifically deplete haematopoietic stem cells (HSC) in adult mice, while leaving peripheral lymphocyte compartments undisturbed. Shortly afterward, the treated mice receive T and B cell-depleted bone marrow from congenic donors, to reconstitute the HSC niche. Donor fractions of 70-90% are established rapidly among HSC, which are stable long-term [23]. Within six weeks of transplant, the fraction of cells that are donor-derived (the chimerism) equilibrates at similar levels at all stages of thymic development and at the transitional stage of B cell development, indicating that turnover of all these populations is complete within this time frame [23]. Donor-derived cells proceed to gradually replace host-derived cells in all peripheral lymphocyte compartments over timescales of weeks to months, first within naive populations and then more gradually within antigen-experience subsets [5]. Importantly, total cell numbers within each subset remained indistinguishable from untreated controls at all timepoints [5], indicating that cells originating from the sex- and age-matched donor and host HSC behaved identically. By using mathematical models to describe the timecourse of replacement within a particular population, one can obtain a direct estimate of the rate of influx of new cells, which can be fed directly into models of DNA labelling [5]. A conceptually similar approach to measuring influx is to use division-linked labelling to explicitly model the kinetics of label in both the population of interest and its precursor. This method is particularly useful when labelling is inefficient, because labelling among precursors is not saturated and is time-varying. Their influx then leaves an informative imprint on any labelling within the target population itself, allowing the rate of ingress of new cells to be inferred [29].

**Fig 1.**
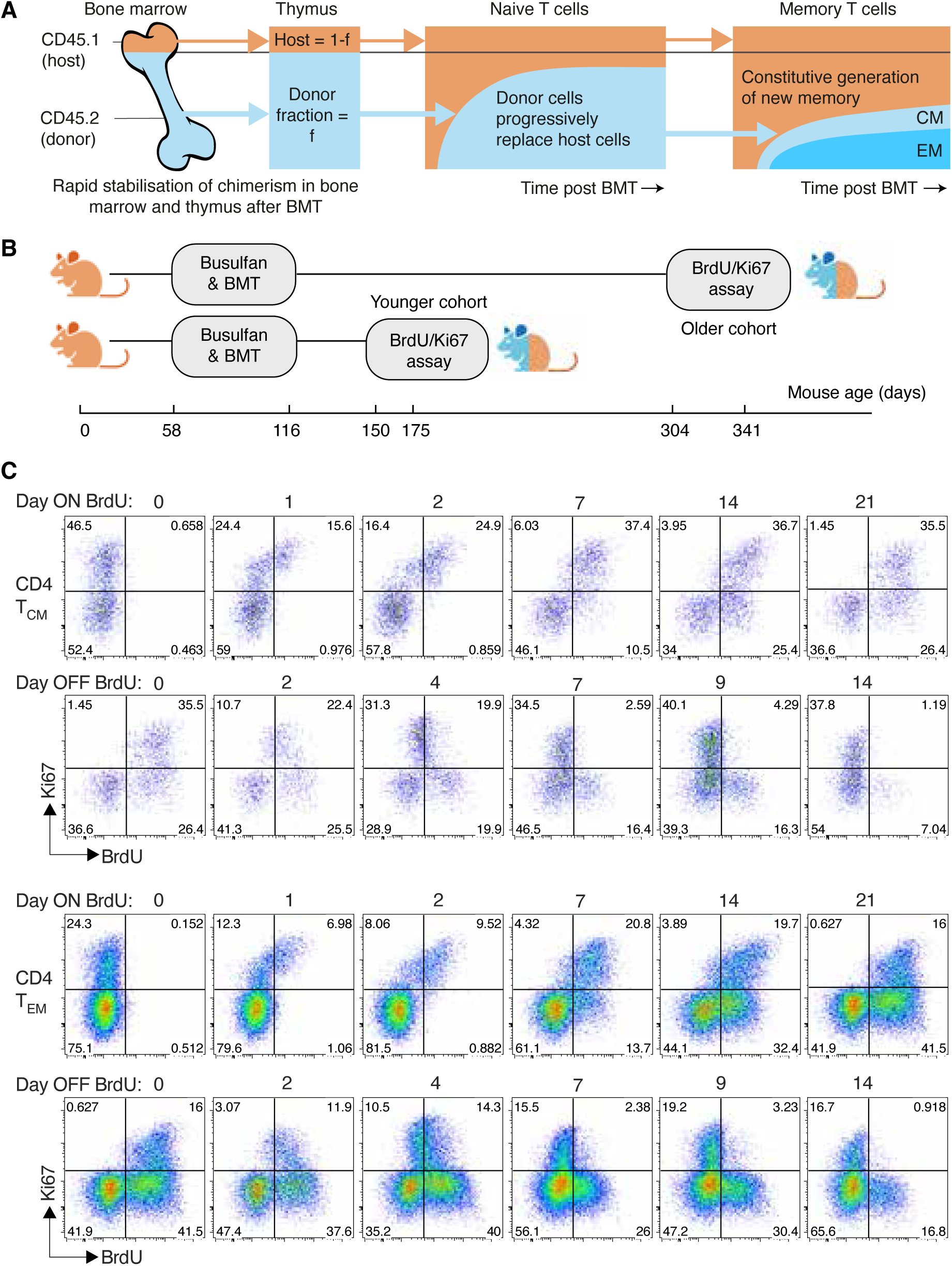
Busulfan chimeric mice, experimental design and gating strategies. **A** Schematic of the busulfan chimeric mouse system, adapted from ref. 32. **B** Design of the BrdU/Ki67 labelling assay, studying two cohorts at different times post BMT. **C** Chimeric mice were fed BrdU for 21 days. Mice were analysed at different times during feeding, and across 14 days following its cessation. Here we show representative timecourses obtained from donor-derived CD4 T_CM_ and T_EM_, showing patterns of Ki67/BrdU expression during and after BrdU feedin5g. See Methods and S1 Fig for details and gating strategies.

Another means of boosting the resolving power of DNA labelling assays is to use concurrent measurements of Ki67, a nuclear protein expressed during mitosis and for a few days afterwards [23, 30, 31]. A cell’s BrdU content and Ki67 expression then report its historical and recent division activity, respectively. This approach was used to study MP CD4 T cell dynamics [5] and rule out a model of ‘temporal heterogeneity’, in which CD4 T_CM_ and T_EM_ cells each comprise populations of cells that divide at a uniform rate, but that a cell’s risk of loss varied with its expression of Ki67. The BrdU/Ki67 approach instead supported a model in which CD4 T_EM_ and T_CM_ each comprise at least two kinetically distinct (‘fast’ and ‘slow’) subpopulations, consistent with other studies [4, 16, 17]. These subpopulations were assumed to form independent lineages that branch from a common precursor [5], but a subsequent study using the busulfan chimera system alone found support for an alternative model in which memory cells progress from fast to slow behaviour [18]. The topologies of the kinetic substructures of CD4 T_CM_ and T_EM_ therefore remain uncertain.

Here, we aimed to characterise the dynamics CD4 T_CM_ and T_EM_ in full, by combining these two approaches – performing BrdU/Ki67 labelling within busulfan chimeric mice directly. Our reasoning was threefold. First, if one defines a memory cell’s age as the time since it or its ancestor entered the memory pool, the average age of donor (newly recruited) clones will be lower than that of the host-derived clones that they are replacing. By stratifying the BrdU/Ki67 analyses by donor and host cells one can therefore quantify any changes in the kinetics of memory T cell clones with their age. Second, by performing the labelling experiments in young and aged cohorts of mice, one can simultaneously observe any changes in dynamics due to the mouse’s chronological age. Third, we hoped that this combined approach would allow us to define the relationship between fast and slow memory CD4 T subsets, and identify the immediate precursors of MP CD4 T_CM_ and T_EM_.

## Results

### Performing DNA labelling assays in busulfan chimeric mice

We studied two age classes of mice (Fig 1B). In one cohort, mice underwent busulfan treatment and bone marrow transplant (BMT) aged between 58 and 116 days and BrdU/Ki67 labelling was performed 56-91 days later (aged between 150-175 days). For brevity we refer to these as ‘young’ mice. An ‘old’ cohort underwent treatment and BMT at similar ages, but with labelling performed between 246 and 260 days later (aged between 304 and 341 days). BrdU was administered by intraperitoneal injection and subsequently in drinking water for up to 21 days. Mice were sacrificed at a range of time points within this period, or up to 14 days after BrdU administration stopped. CD4 T_CM_ and T_EM_ were recovered from lymph nodes, enumerated, and stratified by their BrdU content and Ki67 expression (Fig 1C and S1 Fig).

### The proliferative activity of CD4 T_CM_ and T_EM_ declines as they age

We first analyzed the bulk properties of CD4 MP T cells within the two cohorts. The total numbers of CD4 T_CM_ and T_EM_ recovered from lymph nodes increased slightly with age (Fig 2A). Proliferative activity, measured by the proportion of cells expressing Ki67 (Fig 2B), fell slightly with age among CD4 T_CM_ (median Ki67^high^ fraction = 0.44 in young cohort, 0.36 in older cohort, *p* =0.02) but not significantly for T_EM_ (median Ki67^high^ fraction = 0.20 in both age groups, *p* =0.48).

**Fig 2.**
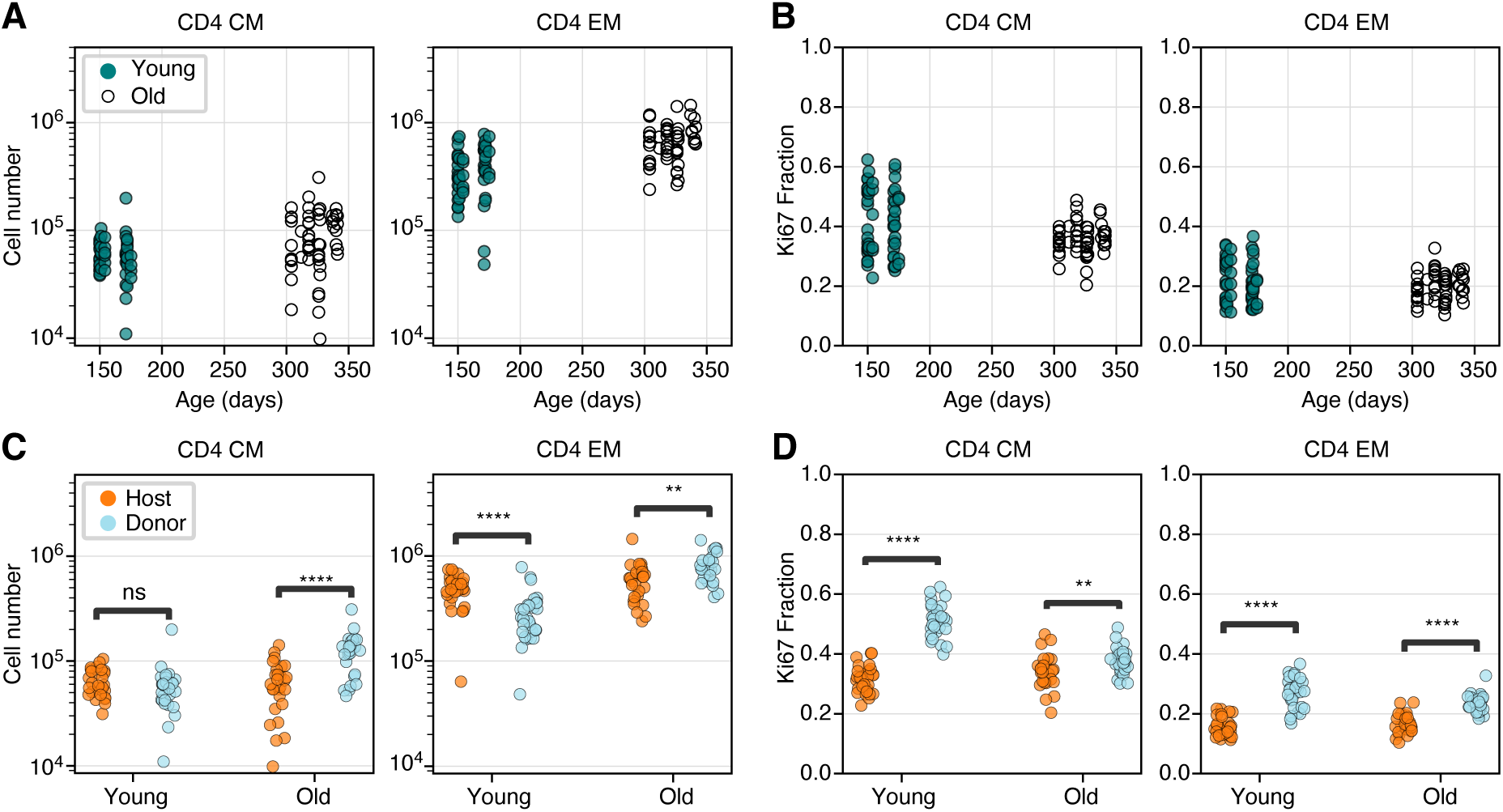
Busulfan chimeric mice reveal shifts in memory CD4 T cell dynamics with both mouse age and cell age. **A, B:** Numbers and Ki67 expression levels of CD4 T_CM_ and T_EM_ recovered from lymph nodes in young and old cohort (Mann-Whitney tests). **C, D:** Numbers and Ki67 expression levels of CD4 T_CM_ and T_EM_ in young and old mice, stratified by average cell age (host and donor; Wilcoxon tests). The data underlying the graphs shown in the figure can be found in S1 Data. ** *p <* 10*^−^*^3^; *** *p <* 10*^−^*^4^; **** *p <* 10*^−^*^5^.

Consistent with reports of the production of new MP CD4 T cells throughout life [5, 18], and the progressive enrichment of all T cell subsets in donor-derived cells after BMT, the composition of both memory subsets shifted towards donor cells in older mice (Fig 2C). This shift was more marked for T_CM_ (*p <* 10*^−^*^4^) but was also significant for T_EM_ (*p <* 10*^−^*^3^). Strikingly, in the younger cohort, donor T_CM_and T_EM_ both expressed Ki67 at substantially higher frequencies than their host counterparts (Fig 2D; *p <* 10*^−^*^8^ in both cases). These differences diminished with time post BMT but remained significant in the older cohort (*p <* 10*^−^*^3^ for both T_CM_ and T_EM_).

These patterns of donor/host cell differences indicate that the average levels of proliferation within cohorts of CD4 T_CM_ and T_EM_ decline with the time since they or their ancestors entered the population. This effect is most apparent among T_CM_ but still appreciable for T_EM_.

### Modelling BrdU/Ki67 labelling kinetics in busulfan chimeric mice

To find a mechanistic explanation of these patterns, we then analysed the BrdU/Ki67 timecourses derived from the young and old mice. As described in Methods, and illustrated in Fig 1B, in each age cohort we followed the uptake and loss of BrdU within CD4 T_CM_ and T_EM_, stratified by host and donor cells and by Ki67 expression, during 21 day pulse and 14 day chase periods. This strategy allowed us to isolate the dynamics of new and old memory cells within young and old mice.

We attempted to describe these data using three classes of mathematical model, shown schematically in Fig 3A. We employed two versions of a kinetic heterogeneity model; branched, in which newly generated T_CM_ and T_EM_ both immediately bifurcate into independent subpopulations (A and B) with distinct rates of division and death [5], and linear, in which cells enter memory with phenotype A and mature into a kinetically distinct phenotype B [18] (Fig 3A, models (ii) and (iii)). We also explored a ‘burst’ model, previously described as ‘temporal heterogeneity’ [22], in which each memory subset (T_CM_/T_EM_, host/donor) comprises quiescent cells of type A which are relatively long-lived and when triggered to divide enter a more dynamic state B, undergoing faster divisions with a reduced lifespan. These cells return to the quiescent state at some unknown rate. The burst model is a generalisation of the model we rejected previously [5], in which Ki67^high^ cells, which are actively dividing or divided recently, are lost at a different rate to that of more quiescent Ki67^low^ cells. The burst model is also conceptually similar to that explored in ref. 33, in which a cell’s risk of death changes with the time since its last division. In all our models, new cells entering memory are assumed to be Ki67^high^, reflecting the assumption that they have recently undergone clonal expansion.

**Fig 3.**
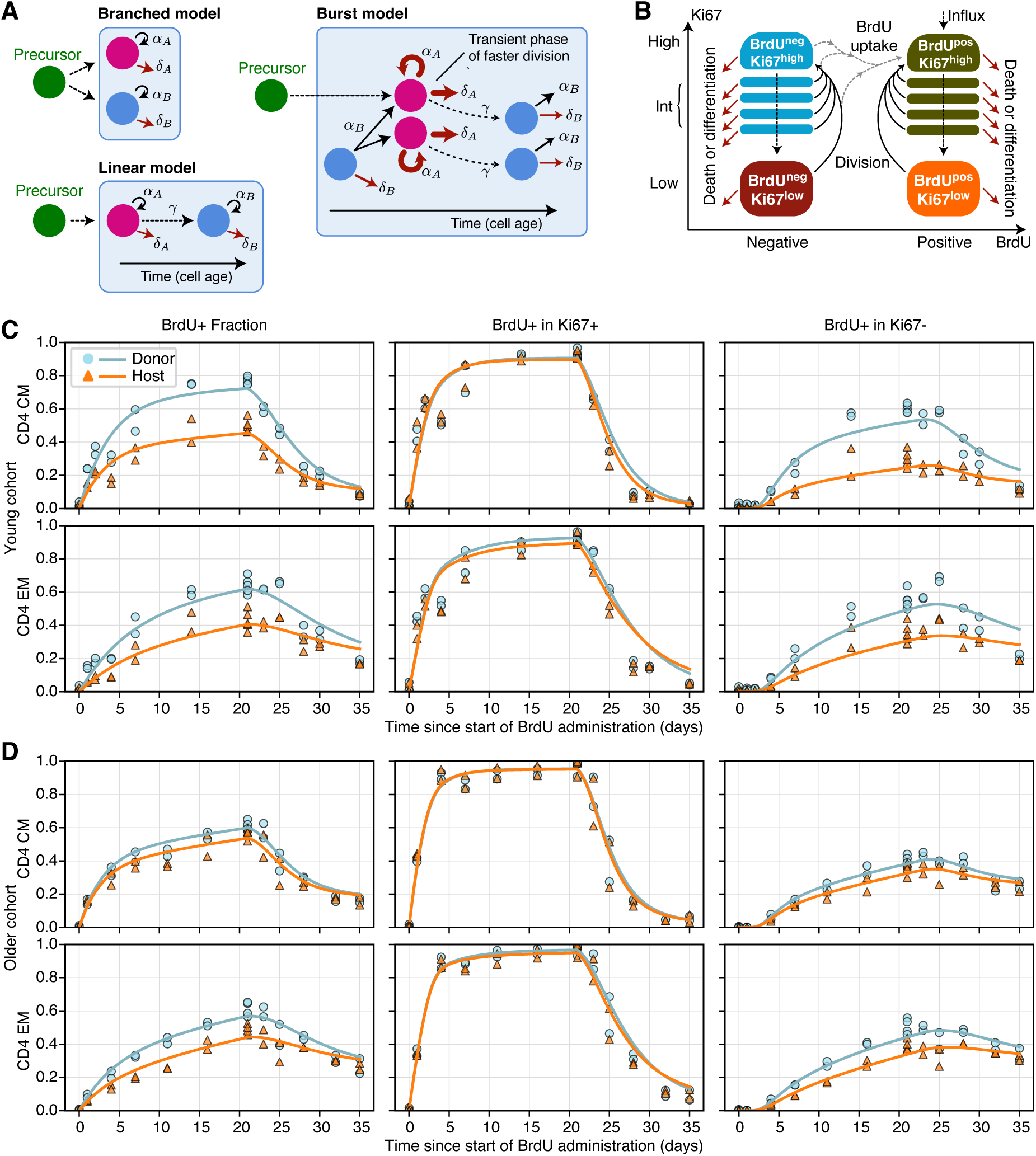
Modelling BrdU/Ki67 timecourses. **A** Models of heterogeneity in cellular dynamics within a memory subset. Adapted from ref. [32]. **B** Schematic of the core component of the ODE model used to describe the uptake of BrdU and the dynamic of Ki67 expression. **C-D** Fitted kinetics of BrdU uptake and loss in CD4 T_CM_ and T_EM_, in the young (**C**) and older (**D**) cohorts of mice, using the branched model. The left column shows the frequency of BrdU-positive cells within donor and host cells. Centre and left columns show the proportion of BrdU-positive cells within Ki67-positive and Ki67-negative cells respectively. There were two mice per time point for both donor and host throughout; in some panels these points are very close, and overlaid. The data underlying the graphs shown in the figure can be found in S1 Data.

We encoded these models with systems of ordinary differential equations. The core structure, which describes the flows between BrdU*^±^* and Ki67^high/low^ cells as they divide and die during the labelling period, is illustrated in Fig 3B. A detailed description of the formulation of the models is in Text A of Supporting Information.

Within both cohorts of mice, the total numbers of host and donor CD4 T_CM_ and T_EM_ cells, and the frequencies of Ki67^high^ cells within each, were approximately constant over the course of the labelling assays (S2 Fig). We therefore made the simplifying assumption that both donor and host memory populations were in quasi-equilibrium, with any changes in their numbers or dynamics occurring over timescales much longer than the 35 day labelling experiment. We took a Bayesian approach to estimating model parameters and performing model selection, using Python and Stan, which is detailed in Text B of Supporting Information. The code and data needed to reproduce our analyses and figures can be found at github.com/elisebullock/PLOS2024.

### Multiple models of memory CD4 T cell dynamics can explain BrdU/Ki67 labelling kinetics

The BrdU labelling data are summarised in Figs 3C and D, shown as the accumulation and loss of BrdU within each population as a whole (left hand panels) and within Ki67^high^ and Ki67^low^ cells (centre and right panels). In the young cohort (Fig 3C), the labelling kinetics of more established (host) and more recently generated (donor) CD4 T_CM_ and T_EM_ were quite distinct, with slower BrdU uptake within host-derived populations. In addition, heterogeneity was apparent in all labelling timecourses. For a kinetically heterogeneous population, the slope of this curve then declines as the faster-dividing populations become saturated with label, and the majority of new uptake occurs within cells that divide more slowly. Such behaviour was apparent, consistent with the known variation in rates of division and/or loss within both T_CM_ and T_EM_. In the older cohort (Fig 3D) BrdU uptake was slower, differences between donor and host kinetics were less apparent, and the labelling kinetics of host-derived cells were similar to those in the younger cohort.

Intuitively, the initial rate of increase of the BrdU-labelled fraction in a population will depend both on the mean division rate of the population and the efficiency of BrdU incorporation per division [34]. In general this efficiency, which we denote *ϵ*, is unknown and this uncertainty confounds the estimation of other parameters [34, 35]. In this regard, the timecourses of the proportion of Ki67^high^ cells that are BrdU-positive (Fig 3C and D, centre panels) are highly informative. Any deviation from a rapid increase to 100% places bounds on *ϵ*, and as we see below, the posterior distributions of this parameter were narrow. Similarly, the kinetics of the BrdU-positive fraction within Ki67^low^ cells are highly informative regarding the lifetime of Ki67 expression – we expect this lifetime to be the delay before the first BrdU-positive Ki67^low^ cells appear, which is clearly apparent in the data (Figs 3C and D, right hand panels) and was again tightly constrained in the model fitting.

To test whether all the parameters of the models illustrated in Fig 3A were identifiable, we fitted each to simulated BrdU/Ki67 labelling timecourses that were generated by the model with added noise. For all models, the estimated total rate of influx of new cells into memory was strongly correlated with the net loss rate of the direct descendant(s) of the source, indicating that these parameters are not separately identifiable. To resolve this, we obtained independent estimates of the rates of influx by modelling the slow accumulation of donor CD4 T_CM_ and T_EM_ within cohorts of busulfan chimeric mice over long timescales (Text C of Supporting Information, and S3 Fig). The posterior distributions of these estimates (summarised in S1 Table) yielded strong priors on the influx rates for the subsequent analyses.

We fitted the branched, linear and burst models to each of the eight timecourses – CD4 T_CM_ and T_EM_separately for donor and host, in both young and old cohorts. All three models yielded excellent fits to the data; surprisingly, these fits were statistically and visually indistinguishable (S2 Table). (Confirming our earlier conclusions [5], the simple temporal heterogeneity model in which Ki67^high^ and Ki67^low^ cells had different death and division rates performed poorly (S4 Fig) and we don’t consider it further here). However, the parameters estimated for the linear and bursting model exhibited some co-linearity (S5 Fig, panels A and B), so we proceed by describing the branched model, whose parameters were most clearly identifiable (S5 Fig, panel C). The fits of the branched model are overlaid in Fig 3C. The posterior distributions of its parameters are shown in Fig 4 and summarised in S3 Table. Corresponding results for the linear and burst models are shown in S6 Fig, S7 Fig and S4 Table, and S8 Fig, S9 Fig and S5 Table respectively. Importantly, however, the conclusions we draw from here onwards are supported by all three models.

**Fig 4.**
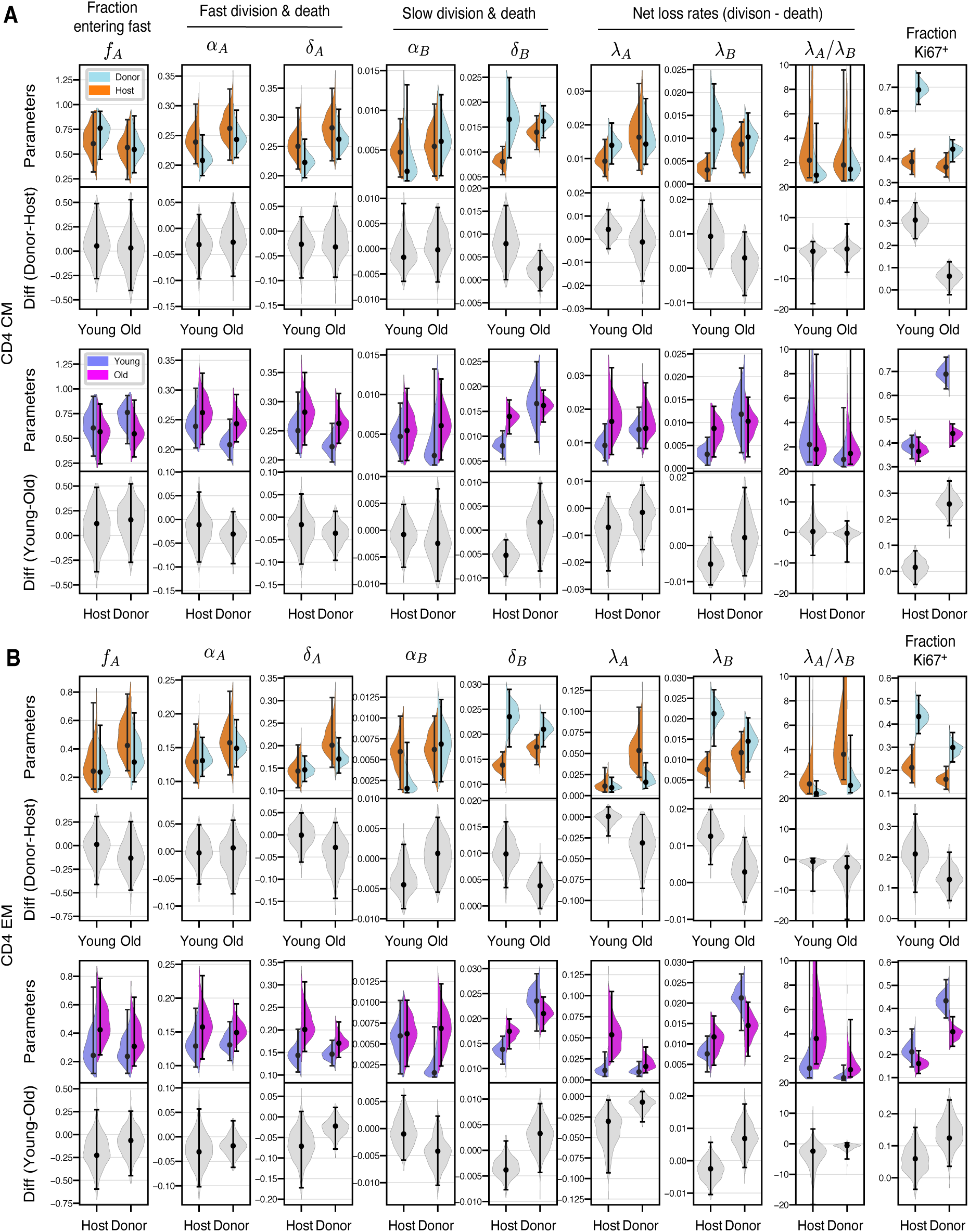
Posterior distributions of parameters derived from the branched model of the kinetics of CD4 T_CM_ (**A**) and T_EM_ (**B**), in the young and old cohorts of mice. Grey violin plots show differences in parameters (host/donor, and young/old). Points and bars show MAP estimates and 95% credible intervals, respectively. The data underlying the graphs shown in the figure can be found in S1 Data.

### Clonal age effects explain changes in CD4 T_CM_ and T_EM_ dynamics with mouse age

For both CD4 T_CM_ and T_EM_, the branched model fits indicated that in both young and older mice, cells that enter the faster subpopulation from the precursor pool divided and were lost over timescales of a few days. In contrast, slow cells divided rarely (mean interdivision times of 100-200 days) and had lifespans of 60-100 days (Fig 4). While the differences in these two subpopulations are clear, these division and loss rate estimates are rather broad because they still exhibited some co-linearity (S5 Fig, panel C). We return to the issue of estimating memory cell lifetimes below. Quantities that were better defined, and perhaps immunologically more relevant, were the balance of loss (at rate *δ*) and self-renewal (at rate *α*) for each subpopulation. This net loss rate, *λ* = *δ −α*, defines the loss of a population in bulk, rather than the turnover of its constituent cells. For example, if the rate at which cells are lost is is balanced exactly by their rate of self-renewal, *λ* = 0 and the population is sustained indefinitely (that is, it is in dynamic equilibrium). In the context of our study, the time taken for a cohort of cells entering a population to halve in size is then ln 2/*λ*. We estimated this quantity for the T_CM_ and T_EM_ populations in bulk, but this half-life is particularly relevant because it simultaneously defines the persistence of a TCR clone. To emphasise this point we refer to ln 2/*λ* as a ‘clonal’ half-life, even though we do not track individual clonotypes in our study. The similarity in the rates of division and loss of fast cells implied that fast CD4 T_CM_ and T_EM_ populations were remarkably persistent, with clonal half lives of between 35-70 days, and T_EM_ slightly less persistent than T_CM_. In contrast, the slower memory cells self-renewed rarely, and so their clonal half-lives (70-140 days) were governed largely by cell lifespan, and were similar for T_CM_ and T_EM_.

Going beyond these broad characterisations, we wanted to identify the basis of the observed host/donor differences, and any changes in memory dynamics with cell age or mouse age. In particular, in young mice, for both CD4 T_CM_ and T_EM_, we inferred that the higher expression of Ki67 within donor cells was due to a greater prevalence of fast cells, compared to host (Fig 4). In Text D of Supporting Information we show that these differences arise because slow, host-derived memory cells were more persistent than their younger donor counterparts, while fast cells behaved similarly, irrespective of cell age (host/donor status) or mouse age. We also found that this increase in the persistence of slow memory clones with their age derived from increased cell lifespan, rather than an increase in self renewal. These conclusions held for all three (branched, linear and burst) models.

We expected the age structure of host-derived CD4 T_CM_ and T_EM_ to change between young and old cohorts to a lesser extent than that of the donor cells. Therefore, any changes linked directly to mouse age ought to be more manifest among host cells. However we saw no shifts in host cell kinetics between the age cohorts, reflected in the similarity of their BrdU/Ki67 labelling curves (Fig 3, panels C and D). Therefore we could not detect any effects of mouse age on CD4 memory dynamics. We can also conclude that increases in the survival capacity of slow CD4 T_CM_ and T_EM_ with cell age must saturate over a timescale of weeks; if they did not, we would expect to see a decline in the loss rates of host-derived memory cells in older mice.

In summary, the stark differences in donor and host kinetics in younger mice are readouts of the effects of cell age, defined as the time since a cell or its ancestor entered the CD4 T_CM_ or T_EM_ population. As they age, slow CD4 T_CM_ and T_EM_ clones become more persistent, shifting the population as a whole to a more quiescent state and explaining the decline in Ki67 within these subsets as mice age. The convergent dynamics of host and donor cells in older mice reflect a convergence in their cell age profiles.

### Comparing memory CD4 T cell lifespans to previous estimates

The other DNA labelling studies we are aware of that measured or inferred memory CD4 T cell lifespans in mice reported the population average, and did not distinguish T_CM_ and T_EM_ [4, 8, 34]. For comparison, we used three independent methods of deriving the corresponding mean lifespan from our data.

Westera *et al.* [4], using heavy water labelling, showed that it is necessary to allow for multiple sub-populations with different rates of turnover to obtain parameter estimates that are independent of the labelling period. They were not able to resolve the number of subpopulations or their rates of turnover individually, but established a robust estimate of the mean lifespan of 15 days (range 11-15) in adult mice. Baliu-Pique *et al.* [8] employed the same approach to derive a lifespan of 8 days (95% CI 5-15). To obtain an analogous estimate, we used the model parameters to calculate the average loss rate of fast and slow cells among both donor and host, each weighted by their population sizes (Text E of Supporting Information). Fig 5A shows the posterior distributions of the lifespans of T_CM_ and T_EM_ separately, with point estimates of roughly 8 and 21 days respectively, and with little change with mouse age. Notably, our confidence in these quantities is greater than our confidence in the lifespans of fast and slow populations themselves (Fig 4). To calculate a population-average lifespan, we account for the lower prevalence of T_CM_ relative to T_EM_ (Fig 5B). The resulting weighted estimates were 18 days (95% CrI 16-21) for younger mice, and 20 days (18-22) in the older cohort (Fig 5C).

**Fig 5.**
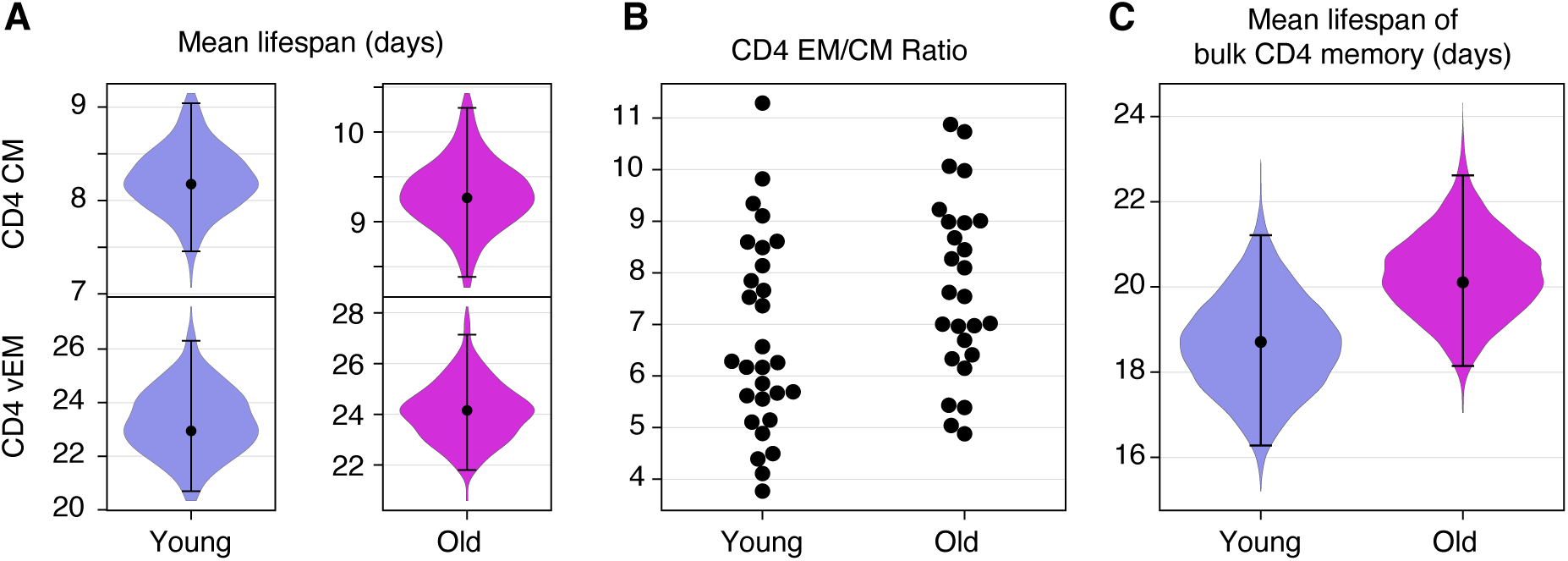
Estimates of the mean lifespan of circulating memory CD4 T cells in the two age cohorts of mice. **A** Expected lifespans of CD4 T_CM_ and T_EM_ in young and old mice, each averaged over fast and slow subpopulations (see Text E of Supporting Information). **B** Relative abundance of lymph node-derived CD4 T_CM_ and T_EM_, by age. **C** Estimated mean lifespans of memory CD4 T cells, averaging over T_CM_ and T_EM_, in the young and old cohorts. Violin plots show the distributions of the lifespans over the joint posterior distribution of all model parameters; also indicated are the median and the 2.5 and 97.5 percentiles. The data underlying the graphs shown in the figure can be found in S1 Data.

We obtained estimates of the mean lifespan of memory CD4 T cells in two other ways. Both rely on estimating the total rate of production, which at steady state balanced by the average loss rate, which is the inverse of the mean lifespan. One method is to use the initial upslope of the BrdU labelling curve, which reflects the rate of production of cells through division and influx. One can show that the mean lifespan is approximately 2*ɛ*/*p*, where *ɛ* is the efficiency of BrdU uptake and *p* is the early rate of increase of the BrdU^+^ fraction (Text F of Supporting Information). In the young cohort, these rates were roughly 0.08/day and 0.04/day for T_CM_ and T_EM_ respectively; in older mice, 0.1 and 0.05. Accounting again for the T_CM_ and T_EM_ abundances (Fig 5B), we obtain rough point estimates of the lifespan of 25 days in the younger cohort and 27 days in the older mice. A similar calculation was employed by De Boer and Perelson [34], who drew on BrdU labelling data from Younes *et al.* [17] to obtain a mean lifespan of between 14 and 22 days (Text F of Supporting Information).

The second method is to utilise Ki67 expression, which again reflects the production of cells through both influx and cell division. If the fraction of cells expressing Ki67 is *k*, and Ki67 is expressed for *T* days after division, one can show that the mean lifespan is approximately *−T* / ln(1 *− k*/2) (Text F of Supporting Information). The frequencies of Ki67 expression within CD4 T_CM_ and T_EM_ were roughly 0.4*±*0.1 and 0.2*±*0.1 respectively, and were similar in young and old mice (Fig 2B); using these values weighted by the abundances of T_CM_ and T_EM_ (Fig 5B, EM/CM *≃* 7.5 *±* 2), with our estimated *T* = 3.1d gives a mean lifespan of approximately 25 days, although with some uncertainty (conservative range 15-70d).

In summary, we find consistent estimates of the average lifespan of memory CD4 T cells in mice that increase slightly with age and are in line with estimates from other studies.

### Using patterns of chimerism to identify the precursors of CD4 T_CM_ and T_EM_

These models of CD4 T_CM_ and T_EM_ dynamics made no assumptions about the identity of their precursors – they only assumed that the rates of influx of host and donor precursors into each subset were constant over the course of the labelling assay, and that new memory cells expressed Ki67 at high levels. Because the donor/host composition of this influx presumably reflects the chimerism of the precursor population, we reasoned that by comparing the chimerism of the flows into CD4 T_CM_ and T_EM_ to that of other populations we might construct a map of the differentiation of CD4 MP T cell subsets.

Fig 6 shows these chimerism estimates. Those inferred from modelling are shown with violin plots (derived from the posterior distributions of parameters in both the branched and linear models), and the points represent experimental observations. Our analysis of the rate of accumulation of donor cells within CD4 T_CM_ and T_EM_ in busulfan chimeric mice (Text C of Supporting Information, and S3 Fig) yielded the chimerism of the flows into these subsets, shown on the left of Fig 6.

**Fig 6.**
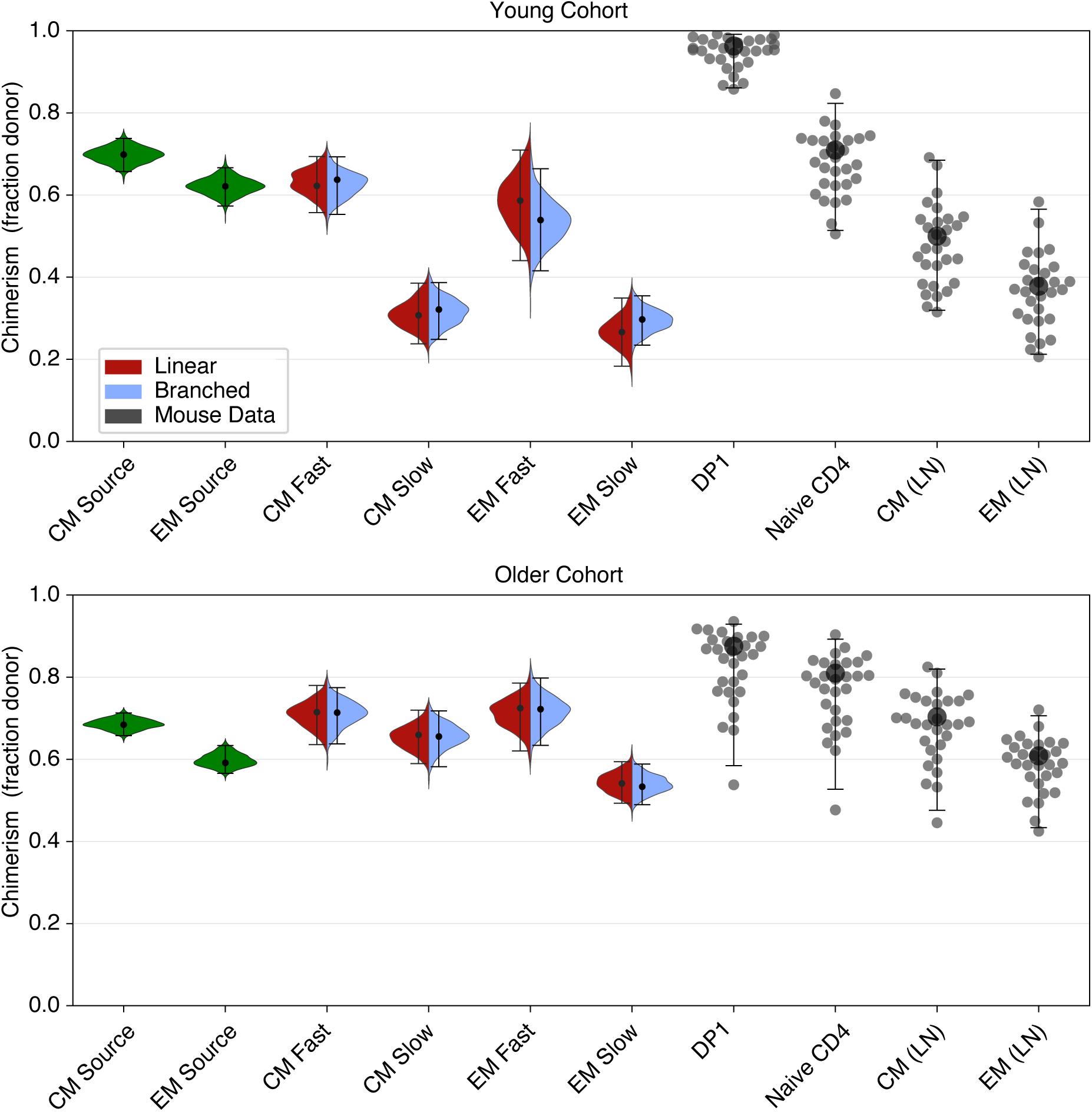
Inferred and measured values of the chimerism (donor cell fraction) within thymic and peripheral T cell subpopulations. Left-hand (green) violin plots indicate the chimerism of the constitutive influxes into CD4 T_CM_ and T_EM_ estimated using Eq 1 of Text C of Supporting Information, *f_d_*. Split-colour violin plots indicate the posterior distributions of the chimerism within fast and slow subsets of CD4 T_CM_ and T_EM_; red and lilac indicate the linear and branched model predictions, respectively. Points represent experimentally observed donor fractions within early-stage double-positive thymocytes (DP1), naive T cells, and CD4 T_CM_ and T_EM_ derived from lymph nodes. The data underlying the graphs shown in the figure can be found in S1 Data.

As expected, chimerism in different cell populations showed considerably more variation in the young cohort, which had undergone BMT relatively recently, than in the older cohort in which donor/host ratios throughout the peripheral T compartments had had more time to approach equilibrium. We therefore focused on the younger cohort to establish potential differentiation pathways. We also saw very little difference in the parameter estimates derived from the branched and linear models, indicating that our inferences are robust to the choice of model.

One source of new memory might be antigen-specific recent thymic emigrants (RTE), whose chimerism is similar to that of early-stage double positive thymocytes (DP1). However, the chimerism of DP1 cells was much higher than the chimerism of the T_CM_ or T_EM_ precursors, so our analyses do not support RTE as the dominant source of new CD4 MP T cells. Instead, the chimerism of T_CM_ precursors aligned well with that of bulk naive CD4 T cells; and the chimerism of the T_EM_ precursors aligned with that of T_CM_. This pattern clearly suggests a naive *→* T_CM_ *→* T_EM_ developmental pathway for CD4 MP T cells, consistent with the order in which donor-derived cells appear within these subsets in busulfan chimeras (S3 Fig). The modelling also provided a breakdown of the observed chimerism of CD4 T_CM_ and T_EM_ into the fast and slow components. It was clear that the estimated chimerism within the slow cells was too low for them to be an exclusive source for any other population, consistent with their being predominantly a terminal differentiation stage in memory T cell development. Instead, any CD4 T_CM_ feeding the T_EM_ pool are likely derived largely from the faster-dividing T_CM_ subpopulation.

### Age-dependent survival rates explain shortfalls in replacement within T_CM_ and T_EM_

In older busulfan chimeric mice, the replacement curves for CD4 T_CM_ and T_EM_ saturated at donor fractions of approximately 0.7 and 0.6 respectively (S3 Fig and Fig 6), lower than that of naive CD4 T cells (Fig 6). Given the average lifespans of CD4 T_CM_ and T_EM_ of a few weeks, it is then perhaps puzzling that the donor fractions within all three compartments do not equalise at late times after BMT. Our analysis suggests that these shortfalls are an effect of cell-age-dependent survival. The mean lifespan is a rather crude measure of memory turnover, as lifespans are significantly right-skewed; in addition, slower cells increase their life expectancies as they age. This increase will lead to a first-in, last-out effect in which donor T cell clones will always be at a survival disadvantage, on average, relative to their older host counterparts. This effect naturally leads to a shortfall in the donor fraction relative to the precursor (putatively, T_CM_ relative to naive; and T_EM_ relative to T_CM_). As the mice age and the donor and host cell age distributions converge, this shortfall decreases, but still remains (Fig 6).

### Validations of model assumptions and fits in other systems

Previously we showed that the total numbers of memory CD4 T cells are indistinguishable in busulfan chimeric mice and WT controls [5]. However there remains the possibility that the observed differences in the behaviour of host and donor memory CD4 T cells in busulfan chimeras derives in some way from some the impact of drug treatment and HSC transplant, rather than simply from their different age profiles. We therefore sought to use two independent mouse reporter systems to validate the structure of our models, and the results of the model fitting.

First, we used an adoptive transfer approach to test the assumptions underlying our models. *Ki67^mCherry-CreER^ Rosa26^RcagYFP^* mice express a Ki67-mCherry fusion protein and inducible CreERT from the *Mki67* locus [26] together with a *Rosa26^RcagYFP^* Cre reporter construct (Materials and Methods). Treatment of donor mice with tamoxifen therefore induces YFP in cells expressing Ki67. Three days following treatment, we isolated either bulk CD4^+^ conventional T cells or purified CD4^+^ T_EM_ T cells from the lymph nodes of these reporter mice, and adoptively transferred them into wild-type CD45.1 congenic hosts (S10 FigA). As expected, Ki67 levels within YFP^+^ cells were higher than among YFP*^−^* cells (Fig 7A and S10 Fig B); having divided in the days prior to transfer, the YFP^+^ cells were enriched for faster-dividing populations. We then assessed Ki67 expression among the transferred cells seven days after transfer (Fig 7B and S10 Fig C). We made two key observations. First, we found that Ki67 levels within donor-derived CD4 T_EM_ declined when they were transferred alone, but were preserved following the transfer of CD4 T cells in bulk. This result is consistent with our basic assumption of continued production of new, Ki67^high^ CD4 T_EM_ from a precursor population that is within circulating CD4 T cells. Second, we observed that Ki67 levels among YFP^+^ cells also fell, and converged. The decline is consistent with the assumption of kinetic heterogeneity; if memory cells divided and died at a constant rate, we would expect the level of Ki67 within the YFP^+^ cells to be preserved. Notably, the decline and convergence of Ki67 levels within YFP^+^ and YFP*^−^* cells is predicted by all three models (Text G of Supporting Information).

**Fig 7.**
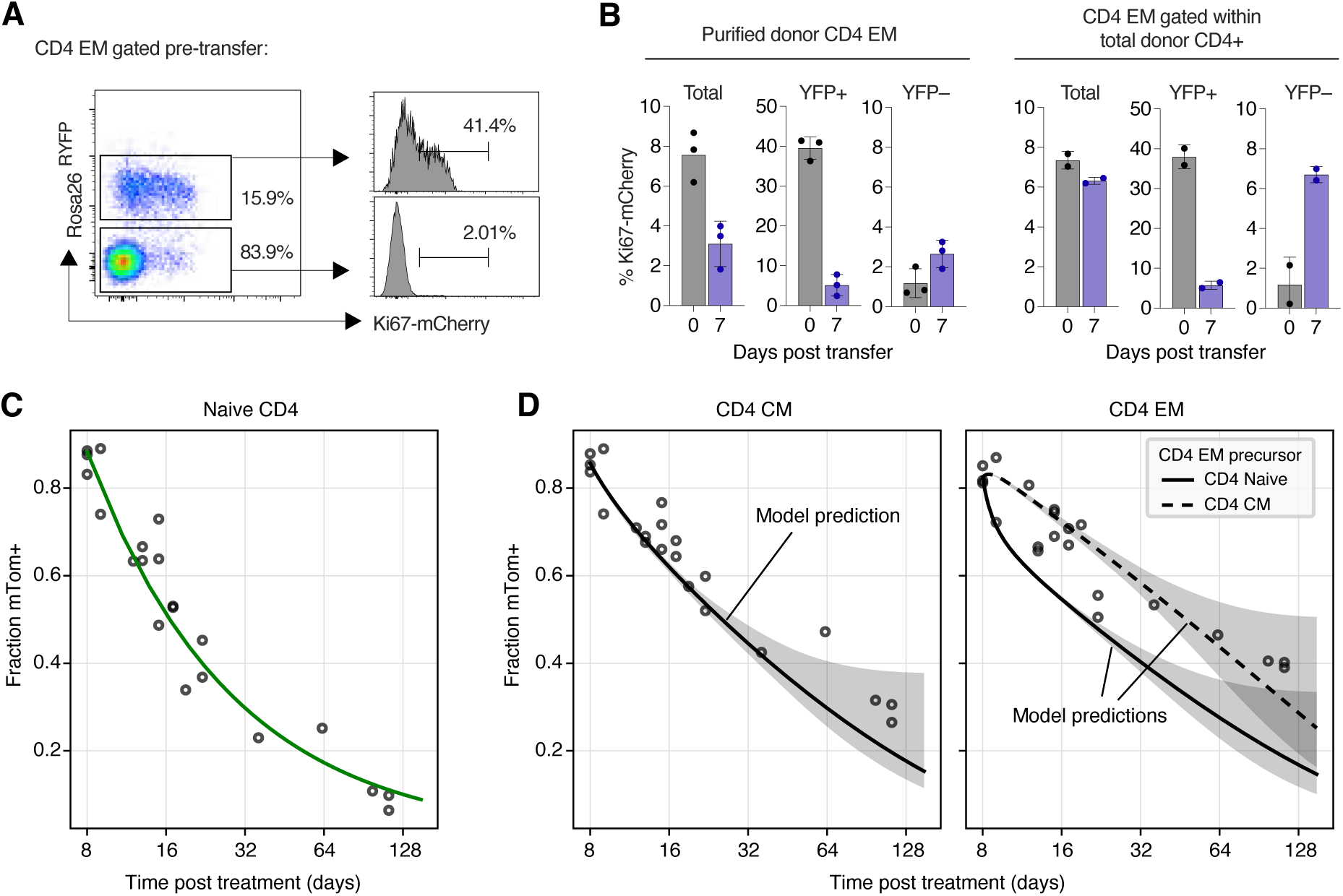
Validation of the structure and predictions of the models of memory CD4 T cell dynamics. **A** CD4 T_EM_ after tamoxifen treatment, pre-transfer, with Ki67 expression stratified by YFP expression. **B** Proportions of cells in bulk, YFP^+^ and YFP*^−^* CD4 T_EM_ expressing Ki67 at days 0 and 7 after transfer. **C** Empirical description (fitted green curve) of the observed time course of the frequency of mTom^+^ cells within the naive CD4 T cell pool, after tamoxifen treatment of CD4 reporter mice. Dilution is driven by the influx of unlabelled cells from the thymus. **D** Observed and predicted frequencies of mTom^+^ cells within the CD4 T_CM_ and T_EM_ following tamoxifen treatment. Shaded regions span the 2.5 and 97.5 percentiles of the distribution of predicted trajectories, generated by sampling 1000 sets of parameters from their joint posterior distribution. The data underlying the graphs shown in the figure can be found in S1 Data.

Second, we used *CD4^CreERT^ Rosa26^RmTom^* mice to directly test the predictions of our fitted models. In this system, the Cre reporter is constructed such that cells expressing CD4 during tamoxifen treatment permanently and heritably express the fluorescent reporter mTomato (mTom). In a closed self-renewing population the frequency of cells expressing mTom will therefore remain constant. Any decline in this frequency therefore reflects the influx of unlabeled precursors. We tamoxifen-treated a cohort of CD4 reporter mice that were age-matched to our younger cohort of busulfan chimeras, and followed them for approximately 18 weeks (S10 FigD). We described the timecourse of the mTom^+^ fraction within naive CD4 T cells empirically (Fig 7C, green curve). We then used the parameters of our fitted models to predict the timecourses of mTom^+^ frequencies within CD4 T_CM_ and T_EM_ (Fig 7D; see Text H of Supporting Information for modelling details). The predictions for both T_CM_ and T_EM_ agreed well with the data, with better visual support for T_CM_ as a precursor to T_EM_ (Fig 7D, rightmost panel). This agreement demonstrates that the kinetics of memory CD4 T cells derived from busulfan chimeric mice are indistinguishable from those in mice that have not undergone HSC depletion and replacement, and so supports our assertion that differences in behavior of host and donor T cells derive only from their different age distributions, and not any ontogenic differences. The simulations also lend support to a dominant CD4 T_CM_ *→* T_EM_ differentiation pathway.

## Discussion

Our key result is that the longer a cohort of CD4 T_CM_ or T_EM_ persists in memory, the greater the expected lifespans of its constituent cells. A consequence of this effect is that the average loss rate of antigen-specific memory CD4 populations decreases as they age. This behaviour mirrors that of naive CD4 and CD8 T cells that we and others have identified in both mice [27, 28, 36] and humans [37], suggesting it is an intrinsic property of T cells.

This dynamic will lead to the accumulation of older memory CD4 T cells. In contrast, a recent study argued that an increase in cell mortality with cell age may act to preserve the TCR diversity of memory populations as an individual ages, and hence be beneficial [38]. In the latter study, cell aging was directly associated with cell divisions, whereas here we defined age to be a heritable clock that measures the time since a cell’s ancestor entered the memory pool. These two measures of age are positively correlated, but any division-aging effects would manifest more strongly among fast memory cells than among slow. Indeed we found that slow memory cells self-renew rarely. It has also been shown that memory cells deriving from aged naive T cells may be functionally impaired [39–41]. It is therefore unclear what cell-age effects are optimal for the long-term preservation of functional memory repertoires.

The mechanistic basis of any change in survival with cell age is unclear. Memory T cells may undergo maturation that increase their expected time-to-die as they age, such as the gradual acquisition of senescence markers. An alternative is a simple selection effect in which every cohort of new memory T cells is heterogeneous in its survival capacity, and more persistent clones are selected for over time. Another possibility is that memory cells experience slow changes in their microenvironments; for example, progressive alterations in receptor expression that might alter their circulation patterns and influence the availability of survival signals. This shift in trafficking patterns would then be a proxy for cell aging. To explore this idea one would need to perform histology to study the spatial distributions of donor- and host-derived memory cells in the younger cohorts of mice, where the average ages of the two populations are most different.

The development and long-term maintenance of memory CD4 T cell subsets has been studied less extensively than those of CD8 T cells, but the dynamics of the latter have some key similarities with the results we describe here, and may help to resolve some of the ambiguities in our analyses. A study involving deuterium labelling of yellow fever virus (YFV)-specific CD8 T cells in humans [42] demonstrated a progressive shift to quiescence over months to years, with surviving memory T cells being very long lived and dividing rarely. Zarnitsyna *et al.* [43] explored a variety of mathematical models to quantify this kinetic, and found strongest support for rates of both division and death that decline with time and result in a power-law decrease in memory cell numbers. Akondy *et al.* [42] found that cells persisting long term were deuterium-rich, indicating that they were derived from populations that divided extensively in the days to weeks following challenge. This kinetic gives support to the linear or burst models of memory rather than the branched models in which the more persistent ‘slow’ cells are quiescent from the moment they enter the memory population.

MP CD4 T cells divide more rapidly on average than conventional antigen-specific memory cells [15–17]. Younes *et al.* [17] argued that MP CD4 T cells comprise a proliferative subset and a slower population with the properties of conventional memory. This distinction was motivated by the observation that bulk and LCMV-specific memory CD4 T cells, identified with tetramer staining 15 days after LCMV challenge, were phenotypically similar, but had Ki67 expression levels of 25-40% and 5% respectively. Bulk memory is presumably enriched for MP cells, and indeed the frequencies of Ki67 expressing cells they observed among the tetramer-negative population are consistent with the roughly 30% weighted average among CD4 T_CM_ and T_EM_ in our analyses (Fig 2B).

There have been conflicting reports regarding the signals required for the generation and maintenance of fast MP CD4 T cells. Younes *et al.* argued that they are driven by cytokines, based on the lack of an effect of an MHC class II-blocking antibody and a modest reduction in proliferation under IL-7R blockade [17]. Their results contrasted with reports that rapidly dividing MP CD4 T cells depend on interactions with MHC class II and require proximal TCR signalling kinases to sustain proliferation [16, 44], and have reduced dependence on IL-7 compared to antigen-specific CD4 memory [15]. However, later studies from the same lab as Younes *et al.* identified the importance of co-stimulation signals from CD28 for the proliferation of fast cells even though division was not affected by MHC class II blockade [12]. The same study highlighted the importance of TCR and CD28 co-stimulation signals for the generation of MP CD4 T cells from naive precursors. These observations can be reconciled if the generation and maintenance of MP CD4 T cells rely on CD28 and TCR interactions but vary in the threshold of TCR signalling required and if there is redundancy with CD28 signals. In addition, interpretation of MHC class II blockade experiments is heavily dependent upon whether antibody blockade is considered absolute, or, as is more likely the case, only partial. If the latter is true, the reduction in availability of MHC class II ligands may be sufficient to block priming events, but insufficient to halt self-renewal of existing memory.

Given the involvement of TCR signals in the production of new MP CD4 T cells, the difference in average levels of proliferation within bulk MP CD4 T cells and antigen-specific memory observed by Younes *et al.* [17] might be explained quite simply by the fact that MP T cells are continually produced in response to chronic stimuli with self or commensal antigens, while LCMV-specific memory will become less proliferative with time post-challenge as the acute infection resolves. The transfer experiment we described here supports this interpretation; CD4 T_EM_ in isolation lost Ki67 expression, whereas when CD4 T cells were transferred in bulk, Ki67 levels on CD4 T_EM_ were maintained, presumably because their precursors were also present and continued to be stimulated. We also see this transition in an infection setting, with levels of Ki67 expression among influenza-specific CD4 T cells declining to low levels 28 days post infection (Jodie Chandler, Ben Seddon, unpublished). Similar reasoning might explain the report by Choo *et al.* that memory CD8 T cells specific for epitopes of LCMV divide at the same rate as memory CD8 T cells in bulk [45]. They established this equivalence by labelling each population with CFSE, transferring them into congenic recipient mice and comparing their CFSE dilution profiles; by doing so, the source of any new, more proliferative MP CD8 cells was presumably removed or diminished, and the remaining polyclonal populations divided at the same slow rate as the LCMV-specific cells. In summary, MP T cells, rather than being distinct from conventional antigen-specific memory, may in fact represent a cross-section of the entire process of memory generation and maintenance.

The binary characterisation of fast and slow that we model here may be a simple representation of a more heterogeneous system. Younes *et al.* [17] saw a 4-fold disparity in Ki67 expression between tetramer-positive (LCMV-specific) and tetramer-negative (MP) CD4 T cells 15 days post-challenge, indicating that fast LCMV-specific memory persisted for only a few days. This transience contrasts to the more extended residence times of fast populations, at the clonal level, that we inferred here. The MP populations we studied likely respond to a variety of commensal or environmental antigens. CD4 T_CM_ and T_EM_ may therefore exhibit variability in their persistence times in a clone-specific way, perhaps depending on antigenic burden, TCR specificity, or their T helper phenotype [46].

By augmenting measurements of BrdU content with Ki67 expression, here and in a previous study we could reject a model of ‘temporal heterogeneity’ in which cells expressing Ki67 are both more likely to divide again and are also at increased risk of loss or onward differentiation. However, the burst model we considered here is a generalisation in which the duration of this more ‘risky’ cell state is no longer tied to expression of Ki67 and is instead a free parameter. We estimated the transition rate from fast to slow to be quite low, such that the average time spent in the bursting state is determined instead by the cell lifespan, which was 4-5 days. Consequently, the efficiency of return to quiescence from the burst phase was rather low, with only *∼* 1% of cells surviving the transition. We found a comparably low efficiency of survival from fast to slow in the linear model. With this additional flexibility, the burst model yielded essentially identical fits to the branched and linear models. We also found that all three models yielded similar predictions for the outcome of the experiment that followed the behaviour of cohorts of cells enriched for fast or slow cells (Fig 7). Therefore the kinetic substructures of CD4 T_CM_ and T_EM_ remain unclear. We speculate they may only be decipherable with single-cell approaches such as barcoding.

Our analyses gave clues as to the sources of new CD4 T_CM_ and T_EM_. By comparing the donor cell content of the constant flow of new cells entering the CD4 T_CM_ and T_EM_ pools to the donor fractions within other cell subsets, we saw clear evidence for a flux from T_CM_ to T_EM_, likely from the rapidly dividing T_CM_ population. We also found evidence that newly produced T_CM_ derive largely from the naive CD4 T cell population in bulk. This was perhaps surprising; among mice housed in clean conditions, without overt infections, we expect naive T cells with appropriate TCR specificities for self or commensal antigens will be recruited into memory efficiently. In that case, one might expect recent thymic emigrants to be the dominant source of new memory, and so for this influx to be highly enriched for donor T cells. Instead, we saw continued recruitment of host cells into memory long after bone marrow transplant, suggesting that even within SPF housing conditions, mice are regularly exposed to new antigenic stimuli. Alternatively, there may be a stochastic component to recruitment, such that naive T cells of any age have the potential to be stimulated to acquire a memory phenotype. Support for the latter idea comes from studies that implicate self antigens as important drivers of conversion to memory [12, 18], and that proliferation is greatest amongst CD5^hi^ cells [12], a marker whose expression correlates with TCR avidity for self. Conversion to memory is therefore likely be an inefficient process since we would expect the naive precursors to be relatively low affinity for self-antigens; the majority of T cells with high affinity for self are deleted in the thymus. The efficiency of recruitment could also be subject to repression by regulatory T cells. Further, as mentioned above, within these relatively low affinity memory populations, any dependence of survival on affinity for self may lead to selection for fitter clones with cell age, and may explain the increase in the average clonal life expectancy over time. The preferential retention of such clones may then be relevant to understanding the increasing incidence of autoimmune disease that occurs with old age. These ideas need to be tested in future studies.

## Material and methods

### Generation of busulfan chimeric mice

Busulfan chimeric C57Bl6/J mice were generated as described in ref. 24. In summary, WT CD45.1 and CD45.2 mice were bred and maintained in conventional pathogen-free colonies at the Royal Free Campus of University College London. CD45.1 mice aged 8 weeks were treated with 20 mg/kg busulfan (Busilvex, Pierre Fabre) to deplete HSC, and reconstituted with T-cell depleted bone marrow cells from congenic donor WT CD45.2 mice. Chimeras were sacrificed at weeks after bone marrow transplantation, Cervical, brachial, axillary, inguinal and mesenteric lymph nodes were dissected from mice; single cell suspensions prepared, and analysed by flow cytometry. BrdU (Sigma) was administered to busulfan chimeric mice by an initial intraperitoneal injection of 0.8mg BrdU, followed by maintenance of 0.8mg/mL BrdU in drinking water for the indicated time periods up to 21 days. BrdU in drinking water was refreshed every 2-3 days. All experiments were performed in accordance with UK Home Office regulations, project license number PP2330953.

### Reporter mouse strains

Ki67^mCherry-CreERT^ reporter mice (*Mki67^tm1.1(cre/ERT2)Bsed^*) have been described previously [26]. These mice were crossed with a Cre reporter strain in which a CAG promoter driven YFP is expressed from the *Rosa26* locus following Cre mediated excision of upstream transcriptional stop sequences; B6.Cg-Gt(ROSA)26^Sortm3(CAG-EYFP)Hze/J^ (Jax strain 007903). Ki67^mCherry-CreERT^ Rosa26^RcagYFP^ mice were homozygous for the indicated mutations at both loci. CD4^CreERT^ Rosa26^RmTom^ reporter mice were generated by breeding Cd4^CreERT2^ mice [47] with B6.Cg-Gt(ROSA)26^Sortm9(CAG-tdTomato)Hze/J^ mice (Jax strain 7909). Experimental mice were heterozygous for the indicated mutations at both loci. Mice were treated with tamoxifen by a single i.p. injection with 2mg of tamxoxifen (Sigma) diluted in corn oil (Fisher Scientific).

### Flow cytometry

Cells were stained with the following monoclonal antibodies and cell dyes: CD45.1 FITC, CD45.2 AlexaFluor 700, TCR-beta APC, CD4 PerCP-eFluor710, CD25 PE, CD44 APC-eFluor780, CD25 eFluor450, CD62L eFluor450 (all eBioscience), TCR-beta PerCP-Cy5.5, CD5 BV510, CD4 BV650, CD44 BV785 (all BioLegend), CD62L BUV737 (BD Biosciences), LIVE/DEAD nearIR and LIVE/DEAD Blue viability dyes (Invitrogen). BrdU and Ki67 co-staining was performed using the FITC BrdU Flow Kit (BD Biosciences) according to the manufacturer’s instructions, along with anti-Ki67 eFluor660 (eBioscience). Cells were acquired on a BD LSR-II or BD LSR-Fortessa flow cytometer and analysed using Flowjo software (Treestar). Subset gates were as follows: CD4 naive: live TCR*β*^+^ CD5^+^ CD4^+^ CD25*^−^* CD44*^−^* CD62L^+^. CD4 T_EM_: live TCR*β*^+^ CD5^+^ CD4^+^ CD25*^−^* CD44^+^ CD62L*^−^*. CD4 T_CM_: live TCR*β*^+^ CD5^+^ CD4^+^ CD25*^−^* CD44^+^ CD62L^+^.

### Cell transfers

For adoptive transfers, donor Ki67^mCherry-CreERT^ Rosa26^RcagYFP^ mice were injected with a single dose of tamoxifen and three days later, total lymph nodes were isolated and CD4 T cells enriched using EasySep Mouse CD4^+^ T cell isolation kit. Cells were then labelled for CD25, CD4, TCR, CD62L and CD44, and total CD4^+^ conventional or CD4 EM further purified by cell sorting on a BD FACS Aria Fusion. Cells (10^6^/mouse) were transferred by tail vein injection into CD45.1 WT hosts. Seven days later, hosts were culled and donor cell phenotype in host lymph nodes analysed by flow.

### Mathematical modelling and statistical analysis

The mathematical models and our approach to model fitting are detailed in Text A and Text B of Supporting Information, respectively. All code and data used to perform model fitting, and details of the prior distributions for parameters, are available at https://github.com/elisebullock/PLOS2024 and also at doi:10.5281/zenodo.11476381. Models were ranked using the Leave-One-Out (LOO) cross validation method [48]. We quantified the relative support for models with the expected log pointwise predictive density (ELPD), for which we report the point estimate with standard error. Models with ELPD differences below 4 were considered indistinguishable.

## Supporting information

**S1 Fig. Complete gating strategy for identifying host and donor CD4 T_CM_ and T_EM_.**

**S2 Fig. Numbers and Ki67 expression of both host and donor CD4 T_CM_ and T_EM_ over the time-courses of the labelling experiments in young and older cohorts of busulfan chimeras.**

**S3 Fig. Estimating the rates of influx into the CD4 T_CM_ and T_EM_ compartments in young and old mice.**

**S4 Fig. Fits of the temporal heterogeneity model to the BrdU/Ki67 labelling data.**

**S5 Fig. Correlations in parameters observed in fits of (A) the linear model, (B) the branched model, and (C) the bursting model.**

**S6 Fig. Fits of the linear model to the BrdU/Ki67 labelling data.**

**S7 Fig. Parameter estimates for the linear model, for CD4 T_CM_ (A) and CD4 T_EM_ (B).**

**S8 Fig. Fits of the bursting model to the BrdU/Ki67 labelling data.**

**S9 Fig. Parameter estimates for the bursting model, for CD4 T_CM_ (A) and CD4 T_EM_ (B).**

**S10 Fig. Fate mapping Ki67-expressing and CD4-expressing cells using reporter mice.**

**S1 Table. MAP estimates and 95% credible intervals on the total daily rate of influx of cells into the CD4 T_CM_ and T_EM_ pools.**

**S2 Table. Comparing support for the three models using the expected log pointwise predictive density (ELPD).**

**S3 Table. MAP estimates and 95% credible intervals on parameters estimated using the branched model.**

**S4 Table. MAP estimates and 95% credible intervals on parameters estimated using the linear model.**

**S5 Table. MAP estimates and 95% credible intervals on parameters estimated using the burst model.**

**S1 Data.** The data underlying the graphs shown in the figures throughout the text and Supporting Information.

**Supporting Information.** Full supporting information, including all supporting figures, tables, and detailed methods. Contains SI Text A-H.

## Acknowledgments

This work was supported by the National Institutes of Health (R01 AI093870 and U01 AI150680) and the Medical Research Council (MR/P011225/1). We are grateful to the reviewers for their insightful critiques.

## Supporting Information

### Supporting Tables

**Table S1.** – MAP estimates and 95% credible intervals on the total daily rate of influx of cells into the CD4 T_CM_ and T_EM_ pools, relative to (Text C, eq. 1; and Fig S3).

| Population | Cohort | Daily influx (MAP and 95% CI) - cells per day |  |
| --- | --- | --- | --- |
|  |  | Donor | Host |
| CD4 T<sub>CM</sub> | Young | 1400 (1200, 1700) | 610 (490, 740) |
|  | Old | 1400 (1200, 1600) | 630 (520, 750) |
| CD4 T<sub>EM</sub> | Young | 14000 (12000, 16000) | 8200 (6900, 9800) |
|  | Old | 13000 (11000, 15000) | 9200 (7500, 11000) |

**Table S2.** – Comparing support for the three models using the expected log pointwise predictive density (ELPD) with standard error (SE).

| Age | Cohort | Population | Model | ELPD | SE |
| --- | --- | --- | --- | --- | --- |
| Old | Host | CD4 T<sub>CM</sub> | Linear | 0 | N/A |
|  |  |  | Branched | 0.25 | 0.78 |
|  |  |  | Burst | 3.86 | 4 |
| Old | Host | CD4 T<sub>EM</sub> | Branched | 0 | N/A |
|  |  |  | Linear | 0.27 | 0.9 |
|  |  |  | Burst | 3.81 | 4.39 |
| Old | Donor | CD4 T<sub>CM</sub> | Linear | 0 | N/A |
|  |  |  | Branched | 0.43 | 0.62 |
|  |  |  | Burst | 7.35 | 5.35 |
| Old | Donor | CD4 T<sub>EM</sub> | Branched | 0 | N/A |
|  |  |  | Linear | 2.34 | 0.85 |
|  |  |  | Burst | 3.1 | 5.66 |
| Young | Host | CD4 T<sub>CM</sub> | Burst | 0 | N/A |
|  |  |  | Linear | 7.44 | 6.2 |
|  |  |  | Branched | 7.89 | 5.99 |
| Young | Host | CD4 T<sub>EM</sub> | Burst | 0 | 0N/A |
|  |  |  | Branched | 0.22 | 0.98 |
|  |  |  | Linear | 1.71 | 0.46 |
| Young | Donor | CD4 T<sub>CM</sub> | Burst | 0 | N/A |
|  |  |  | Branched | 8.64 | 7.36 |
|  |  |  | Linear | 9.39 | 7.47 |
| Young | Donor | CD4 T<sub>EM</sub> | Burst | 0 | N/A |
|  |  |  | Linear | 0.16 | 0.17 |
|  |  |  | Branched | 0.18 | 1.36 |

**Table S3.**
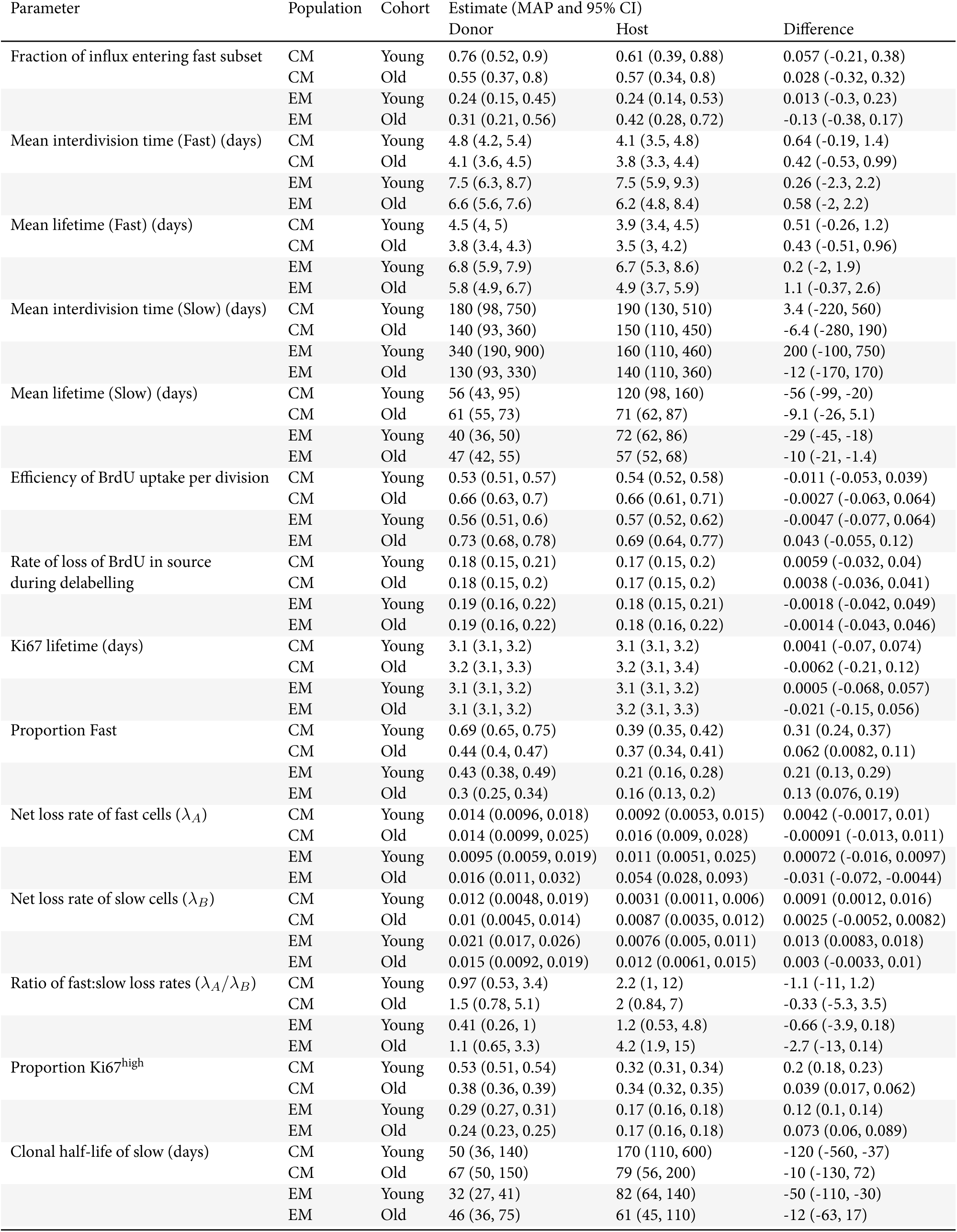
– MAP estimates and 95% credible intervals on parameters estimated using the branched model.

| Parameter | Population | Cohort | Estimate (MAP and 95% CI) |  | Difference |
| --- | --- | --- | --- | --- | --- |
|  |  |  | Donor | Host |  |
| Fraction of influx entering fast subset | CM | Young | 0.76 (0.52, 0.9) | 0.61 (0.39, 0.88) | 0.057 (-0.21, 0.38) |
|  | CM | Old | 0.55 (0.37, 0.8) | 0.57 (0.34, 0.8) | 0.028 (-0.32, 0.32) |
|  | EM | Young | 0.24 (0.15, 0.45) | 0.24 (0.14, 0.53) | 0.013 (-0.3, 0.23) |
|  | EM | Old | 0.31 (0.21, 0.56) | 0.42 (0.28, 0.72) | -0.13 (-0.38, 0.17) |
| Mean interdivision time (Fast) (days) | CM | Young | 4.8 (4.2, 5.4) | 4.1 (3.5, 4.8) | 0.64 (-0.19, 1.4) |
|  | CM | Old | 4.1 (3.6, 4.5) | 3.8 (3.3, 4.4) | 0.42 (-0.53, 0.99) |
|  | EM | Young | 7.5 (6.3, 8.7) | 7.5 (5.9, 9.3) | 0.26 (-2.3, 2.2) |
|  | EM | Old | 6.6 (5.6, 7.6) | 6.2 (4.8, 8.4) | 0.58 (-2, 2.2) |
| Mean lifetime (Fast) (days) | CM | Young | 4.5 (4, 5) | 3.9 (3.4, 4.5) | 0.51 (-0.26, 1.2) |
|  | CM | Old | 3.8 (3.4, 4.3) | 3.5 (3, 4.2) | 0.43 (-0.51, 0.96) |
|  | EM | Young | 6.8 (5.9, 7.9) | 6.7 (5.3, 8.6) | 0.2 (-2, 1.9) |
|  | EM | Old | 5.8 (4.9, 6.7) | 4.9 (3.7, 5.9) | 1.1 (-0.37, 2.6) |
| Mean interdivision time (Slow) (days) | CM | Young | 180 (98, 750) | 190 (130, 510) | 3.4 (-220, 560) |
|  | CM | Old | 140 (93, 360) | 150 (110, 450) | -6.4 (-280, 190) |
|  | EM | Young | 340 (190, 900) | 160 (110, 460) | 200 (-100, 750) |
|  | EM | Old | 130 (93, 330) | 140 (110, 360) | -12 (-170, 170) |
| Mean lifetime (Slow) (days) | CM | Young | 56 (43, 95) | 120 (98, 160) | -56 (-99, -20) |
|  | CM | Old | 61 (55, 73) | 71 (62, 87) | -9.1 (-26, 5.1) |
|  | EM | Young | 40 (36, 50) | 72 (62, 86) | -29 (-45, -18) |
|  | EM | Old | 47 (42, 55) | 57 (52, 68) | -10 (-21, -1.4) |
| Efficiency of BrdU uptake per division | CM | Young | 0.53 (0.51, 0.57) | 0.54 (0.52, 0.58) | -0.011 (-0.053, 0.039) |
|  | CM | Old | 0.66 (0.63, 0.7) | 0.66 (0.61, 0.71) | -0.0027 (-0.063, 0.064) |
|  | EM | Young | 0.56 (0.51, 0.6) | 0.57 (0.52, 0.62) | -0.0047 (-0.077, 0.064) |
|  | EM | Old | 0.73 (0.68, 0.78) | 0.69 (0.64, 0.77) | 0.043 (-0.055, 0.12) |
| Rate of loss of BrdU in source during delabelling | CM | Young | 0.18 (0.15, 0.21) | 0.17 (0.15, 0.2) | 0.0059 (-0.032, 0.04) |
|  | CM | Old | 0.18 (0.15, 0.2) | 0.17 (0.15, 0.2) | 0.0038 (-0.036, 0.041) |
|  | EM | Young | 0.19 (0.16, 0.22) | 0.18 (0.15, 0.21) | -0.0018 (-0.042, 0.049) |
|  | EM | Old | 0.19 (0.16, 0.22) | 0.18 (0.16, 0.22) | -0.0014 (-0.043, 0.046) |
| Ki67 lifetime (days) | CM | Young | 3.1 (3.1, 3.2) | 3.1 (3.1, 3.2) | 0.0041 (-0.07, 0.074) |
|  | CM | Old | 3.2 (3.1, 3.3) | 3.2 (3.1, 3.4) | -0.0062 (-0.21, 0.12) |
|  | EM | Young | 3.1 (3.1, 3.2) | 3.1 (3.1, 3.2) | 0.0005 (-0.068, 0.057) |
|  | EM | Old | 3.1 (3.1, 3.2) | 3.2 (3.1, 3.3) | -0.021 (-0.15, 0.056) |
| Proportion Fast | CM | Young | 0.69 (0.65, 0.75) | 0.39 (0.35, 0.42) | 0.31 (0.24, 0.37) |
|  | CM | Old | 0.44 (0.4, 0.47) | 0.37 (0.34, 0.41) | 0.062 (0.0082, 0.11) |
|  | EM | Young | 0.43 (0.38, 0.49) | 0.21 (0.16, 0.28) | 0.21 (0.13, 0.29) |
|  | EM | Old | 0.3 (0.25, 0.34) | 0.16 (0.13, 0.2) | 0.13 (0.076, 0.19) |
| Net loss rate of fast cells ($\lambda_A$) | CM | Young | 0.014 (0.0096, 0.018) | 0.0092 (0.0053, 0.015) | 0.0042 (-0.0017, 0.01) |
|  | CM | Old | 0.014 (0.0099, 0.025) | 0.016 (0.009, 0.028) | -0.00091 (-0.013, 0.011) |
|  | EM | Young | 0.0095 (0.0059, 0.019) | 0.011 (0.0051, 0.025) | 0.00072 (-0.016, 0.0097) |
|  | EM | Old | 0.016 (0.011, 0.032) | 0.054 (0.028, 0.093) | -0.031 (-0.072, -0.0044) |
| Net loss rate of slow cells ($\lambda_B$) | CM | Young | 0.012 (0.0048, 0.019) | 0.0031 (0.0011, 0.006) | 0.0091 (0.0012, 0.016) |
|  | CM | Old | 0.01 (0.0045, 0.014) | 0.0087 (0.0035, 0.012) | 0.0025 (-0.0052, 0.0082) |
|  | EM | Young | 0.021 (0.017, 0.026) | 0.0076 (0.005, 0.011) | 0.013 (0.0083, 0.018) |
|  | EM | Old | 0.015 (0.0092, 0.019) | 0.012 (0.0061, 0.015) | 0.003 (-0.0033, 0.01) |
| Ratio of fast:slow loss rates ($\lambda_A/\lambda_B$) | CM | Young | 0.97 (0.53, 3.4) | 2.2 (1, 12) | -1.1 (-11, 1.2) |
|  | CM | Old | 1.5 (0.78, 5.1) | 2 (0.84, 7) | -0.33 (-5.3, 3.5) |
|  | EM | Young | 0.41 (0.26, 1) | 1.2 (0.53, 4.8) | -0.66 (-3.9, 0.18) |
|  | EM | Old | 1.1 (0.65, 3.3) | 4.2 (1.9, 15) | -2.7 (-13, 0.14) |
| Proportion Ki67<sup>high</sup> | CM | Young | 0.53 (0.51, 0.54) | 0.32 (0.31, 0.34) | 0.2 (0.18, 0.23) |
|  | CM | Old | 0.38 (0.36, 0.39) | 0.34 (0.32, 0.35) | 0.039 (0.017, 0.062) |
|  | EM | Young | 0.29 (0.27, 0.31) | 0.17 (0.16, 0.18) | 0.12 (0.1, 0.14) |
|  | EM | Old | 0.24 (0.23, 0.25) | 0.17 (0.16, 0.18) | 0.073 (0.06, 0.089) |
| Clonal half-life of slow (days) | CM | Young | 50 (36, 140) | 170 (110, 600) | -120 (-560, -37) |
|  | CM | Old | 67 (50, 150) | 79 (56, 200) | -10 (-130, 72) |
|  | EM | Young | 32 (27, 41) | 82 (64, 140) | -50 (-110, -30) |
|  | EM | Old | 46 (36, 75) | 61 (45, 110) | -12 (-63, 17) |

**Table S4.** – MAP estimates and 95% credible intervals on parameters estimated using the linear model.

| Parameter | Population | Cohort | Estimate (MAP and 95% CI) |  | Difference |
| --- | --- | --- | --- | --- | --- |
|  |  |  | Donor | Host |  |
| Rate of fast to slow | CM | Young | 0.0028 (0.0015, 0.0097) | 0.0041 (0.0018, 0.01) | 0.00033 (-0.0068, 0.0052) |
|  | CM | Old | 0.0056 (0.0019, 0.018) | 0.0048 (0.0018, 0.018) | 0.00078 (-0.012, 0.012) |
|  | EM | Young | 0.0071 (0.0024, 0.028) | 0.0059 (0.0028, 0.028) | 0.00082 (-0.019, 0.021) |
|  | EM | Old | 0.026 (0.01, 0.033) | 0.036 (0.012, 0.053) | -0.01 (-0.035, 0.016) |
| Mean interdivision time(Fast) (days) | CM | Young | 4.9 (4.2, 5.4) | 4.1 (3.5, 4.8) | 0.71 (-0.19, 1.6) |
|  | CM | Old | 4 (3.6, 4.6) | 3.7 (3.2, 4.4) | 0.32 (-0.55, 1.1) |
|  | EM | Young | 7.9 (6.7, 11) | 7.3 (5.8, 10) | 0.51 (-2.6, 3.7) |
|  | EM | Old | 6.3 (5.3, 7.5) | 5.8 (4.7, 7.5) | 0.61 (-1.4, 2) |
| Mean lifetime (Fast) (days) | CM | Young | 4.6 (3.9, 5) | 3.9 (3.4, 4.6) | 0.55 (-0.24, 1.3) |
|  | CM | Old | 3.8 (3.3, 4.3) | 3.4 (3, 4.1) | 0.31 (-0.55, 1) |
|  | EM | Young | 7.8 (6.1, 9.1) | 6.9 (5, 8.6) | 1.6 (-1.7, 3.1) |
|  | EM | Old | 6 (5.1, 7.1) | 5.1 (4.2, 6.7) | 0.64 (-0.9, 2.2) |
| Mean interdivision time (Slow) (days) | CM | Young | 140 (81, 550) | 160 (120, 390) | -20 (-210, 390) |
|  | CM | Old | 97 (78, 220) | 100 (84, 220) | -7.8 (-110, 110) |
|  | EM | Young | 120 (64, 730) | 91 (77, 140) | 20 (-45, 630) |
|  | EM | Old | 94 (74, 140) | 100 (87, 170) | -8 (-69, 36) |
| Mean lifetime (Slow) (days) | CM | Young | 52 (40, 100) | 110 (93, 150) | -55 (-93, -4.8) |
|  | CM | Old | 63 (51, 77) | 74 (60, 95) | -13 (-35, 8.9) |
|  | EM | Young | 46 (36, 110) | 73 (64, 95) | -31 (-49, 29) |
|  | EM | Old | 49 (44, 60) | 62 (55, 77) | -15 (-28, -0.32) |
| Efficiency of BrdU uptake per division | CM | Young | 0.53 (0.51, 0.57) | 0.54 (0.51, 0.59) | -0.0013 (-0.059, 0.038) |
|  | CM | Old | 0.67 (0.63, 0.71) | 0.68 (0.63, 0.73) | -0.011 (-0.069, 0.062) |
|  | EM | Young | 0.58 (0.53, 0.64) | 0.61 (0.56, 0.66) | -0.019 (-0.099, 0.056) |
|  | EM | Old | 0.77 (0.72, 0.81) | 0.76 (0.7, 0.81) | 0.0067 (-0.066, 0.089) |
| Rate of loss of BrdU in source during delabelling | CM | Young | 0.17 (0.15, 0.21) | 0.17 (0.15, 0.2) | 0.0056 (-0.032, 0.043) |
|  | CM | Old | 0.17 (0.15, 0.2) | 0.17 (0.15, 0.2) | 0.0051 (-0.034, 0.042) |
|  | EM | Young | 0.19 (0.16, 0.22) | 0.18 (0.15, 0.21) | -0.0024 (-0.036, 0.046) |
|  | EM | Old | 0.17 (0.15, 0.21) | 0.18 (0.15, 0.21) | -0.0085 (-0.043, 0.039) |
| Ki67 lifetime (days) | CM | Young | 3.1 (3.1, 3.2) | 3.1 (3.1, 3.2) | -0.0034 (-0.087, 0.059) |
|  | CM | Old | 3.1 (3.1, 3.3) | 3.2 (3.1, 3.4) | -0.0091 (-0.25, 0.11) |
|  | EM | Young | 3.1 (3.1, 3.2) | 3.1 (3.1, 3.2) | -0.00099 (-0.066, 0.046) |
|  | EM | Old | 3.1 (3.1, 3.2) | 3.2 (3.1, 3.3) | -0.0098 (-0.14, 0.045) |
| Proportion Fast | CM | Young | 0.72 (0.65, 0.75) | 0.38 (0.35, 0.42) | 0.32 (0.25, 0.38) |
|  | CM | Old | 0.43 (0.4, 0.47) | 0.37 (0.33, 0.41) | 0.067 (0.008, 0.11) |
|  | EM | Young | 0.51 (0.38, 0.62) | 0.21 (0.15, 0.27) | 0.31 (0.15, 0.42) |
|  | EM | Old | 0.29 (0.25, 0.35) | 0.16 (0.14, 0.21) | 0.13 (0.07, 0.19) |
| Net loss rate of fast cells ($\lambda_A$) | CM | Young | 0.013 (0.0085, 0.019) | 0.01 (0.0038, 0.015) | 0.0019 (-0.0036, 0.012) |
|  | CM | Old | 0.019 (0.0067, 0.027) | 0.019 (0.0086, 0.03) | -0.0027 (-0.017, 0.013) |
|  | EM | Young | 0.017 (0.0025, 0.027) | 0.023 (0.0049, 0.041) | -0.011 (-0.03, 0.014) |
|  | EM | Old | 0.003 (0.00047, 0.021) | 0.0092 (0.0027, 0.047) | -0.0051 (-0.042, 0.0092) |
| Net loss rate of slow cells ($\lambda_B$) | CM | Young | 0.014 (0.0038, 0.02) | 0.0025 (0.0011, 0.0065) | 0.011 (0.000059, 0.017) |
|  | CM | Old | 0.0044 (0.0014, 0.014) | 0.0029 (0.00094, 0.011) | 0.0017 (-0.0063, 0.01) |
|  | EM | Young | 0.0078 (0.003, 0.024) | 0.0016 (0.00063, 0.0072) | 0.0051 (-0.0012, 0.022) |
|  | EM | Old | 0.011 (0.0043, 0.014) | 0.0071 (0.0022, 0.011) | 0.0027 (-0.0039, 0.0096) |
| Ratio of fast:slow loss rates ($\lambda_A/\lambda_B$) | CM | Young | 0.88 (0.44, 4.7) | 2.3 (0.64, 12) | -1.1 (-9.7, 2.4) |
|  | CM | Old | 2.1 (0.51, 18) | 3.7 (0.88, 28) | -1.3 (-24, 13) |
|  | EM | Young | 0.87 (0.099, 8.1) | 6.3 (0.7, 58) | -4.7 (-55, 3.4) |
|  | EM | Old | 0.4 (0.034, 4.8) | 2.1 (0.25, 22) | -1.3 (-21, 2.5) |
| Proportion Ki67<sup>high</sup> | CM | Young | 0.53 (0.51, 0.55) | 0.32 (0.3, 0.34) | 0.2 (0.18, 0.23) |
|  | CM | Old | 0.38 (0.36, 0.39) | 0.33 (0.32, 0.36) | 0.037 (0.016, 0.061) |
|  | EM | Young | 0.3 (0.28, 0.32) | 0.17 (0.16, 0.18) | 0.13 (0.11, 0.16) |
|  | EM | Old | 0.24 (0.23, 0.26) | 0.17 (0.16, 0.18) | 0.075 (0.059, 0.089) |
| Clonal half-life of slow (days) | CM | Young | 49 (35, 180) | 180 (110, 620) | -130 (-540, -0.86) |
|  | CM | Old | 91 (49, 490) | 140 (64, 740) | -43 (-610, 290) |
|  | EM | Young | 48 (29, 230) | 210 (97, 1100) | -140 (-1100, 43) |
|  | EM | Old | 64 (50, 160) | 90 (62, 320) | -23 (-250, 56) |

**Table S5.** – MAP estimates and 95% credible intervals on parameters estimated using the burst model.

| Parameter | Population | Cohort | Estimate (MAP and 95% CI) |  | Difference |
| --- | --- | --- | --- | --- | --- |
|  |  |  | Donor | Host |  |
| Rate of fast to slow | CM | Young | 0.0051 (0.0032, 0.0096) | 0.01 (0.0077, 0.016) | -0.0045 (-0.011, 0.00033) |
|  | CM | Old | 0.021 (0.015, 0.027) | 0.024 (0.017, 0.032) | -0.0023 (-0.012, 0.006) |
|  | EM | Young | 0.0097 (0.0027, 0.021) | 0.02 (0.0099, 0.04) | -0.011 (-0.03, 0.0041) |
|  | EM | Old | 0.037 (0.026, 0.05) | 0.063 (0.044, 0.086) | -0.025 (-0.053, -0.0028) |
| Mean interdivision time (Fast) (days) | CM | Young | 4.4 (3.8, 4.7) | 3.8 (3.3, 4.3) | 0.58 (-0.22, 1.1) |
|  | CM | Old | 4 (3.7, 4.6) | 3.9 (3.3, 4.5) | 0.099 (-0.65, 0.92) |
|  | EM | Young | 7.4 (6.3, 8.8) | 7.9 (6.6, 9.7) | -0.24 (-2.4, 1.5) |
|  | EM | Old | 6.6 (5.9, 7.5) | 6.9 (5.4, 8.2) | 0.23 (-1.7, 1.5) |
| Mean lifetime (Fast) (days) | CM | Young | 4.1 (3.6, 4.5) | 3.6 (3.2, 4.1) | 0.52 (-0.29, 0.99) |
|  | CM | Old | 3.8 (3.4, 4.2) | 3.6 (3.1, 4.2) | 0.14 (-0.56, 0.81) |
|  | EM | Young | 6.7 (6, 7.8) | 7.5 (6.2, 9) | -0.61 (-2.3, 1) |
|  | EM | Old | 6.3 (5.5, 7.2) | 6.1 (4.8, 7.3) | 0.31 (-1.2, 1.8) |
| Mean interdivision time (Slow) (days) | CM | Young | 150 (100, 350) | 280 (200, 370) | -130 (-230, 83) |
|  | CM | Old | 120 (98, 170) | 150 (110, 170) | -17 (-53, 34) |
|  | EM | Young | 340 (140, 1300) | 220 (140, 970) | 98 (-480, 1000) |
|  | EM | Old | 120 (88, 220) | 110 (97, 210) | -15 (-78, 90) |
| Mean lifetime (Slow) (days) | CM | Young | 84 (49, 140) | 190 (130, 270) | -100 (-200, -32) |
|  | CM | Old | 88 (68, 120) | 100 (80, 130) | -14 (-54, 22) |
|  | EM | Young | 61 (41, 230) | 96 (77, 190) | -43 (-120, 120) |
|  | EM | Old | 57 (50, 80) | 73 (63, 99) | -15 (-42, 5.8) |
| Efficiency of BrdU uptake per division | CM | Young | 0.51 (0.48, 0.54) | 0.51 (0.48, 0.55) | -0.005 (-0.053, 0.046) |
|  | CM | Old | 0.65 (0.61, 0.68) | 0.65 (0.61, 0.7) | -0.0058 (-0.062, 0.051) |
|  | EM | Young | 0.52 (0.49, 0.57) | 0.5 (0.47, 0.56) | 0.018 (-0.035, 0.077) |
|  | EM | Old | 0.71 (0.67, 0.76) | 0.7 (0.64, 0.75) | 0.031 (-0.056, 0.086) |
| Rate of loss of BrdU in source during delabeling | CM | Young | 0.17 (0.14, 0.2) | 0.16 (0.14, 0.2) | 0.0041 (-0.035, 0.039) |
|  | CM | Old | 0.16 (0.14, 0.19) | 0.16 (0.14, 0.2) | -0.005 (-0.037, 0.037) |
|  | EM | Young | 0.18 (0.15, 0.2) | 0.17 (0.15, 0.21) | 0.0022 (-0.038, 0.043) |
|  | EM | Old | 0.17 (0.15, 0.21) | 0.17 (0.15, 0.2) | 0.0057 (-0.041, 0.04) |
| Ki67 lifetime (days) | CM | Young | 2.3 (2.1, 2.4) | 2.2 (2, 2.4) | 0.052 (-0.23, 0.31) |
|  | CM | Old | 2.7 (2.5, 3) | 2.8 (2.6, 3.1) | -0.071 (-0.47, 0.28) |
|  | EM | Young | 1.9 (1.8, 2.1) | 2 (1.8, 2.1) | -0.0011 (-0.2, 0.19) |
|  | EM | Old | 2.5 (2.3, 2.7) | 2.7 (2.4, 2.9) | -0.16 (-0.48, 0.21) |
| Proportion Fast | CM | Young | 0.76 (0.72, 0.8) | 0.44 (0.41, 0.48) | 0.31 (0.26, 0.37) |
|  | CM | Old | 0.48 (0.45, 0.51) | 0.41 (0.38, 0.45) | 0.052 (0.018, 0.11) |
|  | EM | Young | 0.63 (0.56, 0.72) | 0.38 (0.31, 0.44) | 0.26 (0.15, 0.37) |
|  | EM | Old | 0.4 (0.35, 0.44) | 0.25 (0.21, 0.29) | 0.16 (0.086, 0.21) |
| Net loss rate of fast cells ($\lambda_A$) | CM | Young | 0.014 (0.0092, 0.021) | 0.011 (0.0066, 0.016) | 0.0037 (-0.0039, 0.01) |
|  | CM | Old | 0.018 (0.012, 0.029) | 0.018 (0.0094, 0.031) | 0.00022 (-0.013, 0.014) |
|  | EM | Young | 0.013 (0.0029, 0.019) | 0.008 (0.00087, 0.016) | -0.00024 (-0.0073, 0.014) |
|  | EM | Old | 0.0053 (0.001, 0.02) | 0.011 (0.0027, 0.038) | -0.0057 (-0.031, 0.0086) |
| Net loss rate of slow cells ($\lambda_B$) | CM | Young | 0.0056 (0.00064, 0.016) | 0.00082 (0.00012, 0.0045) | 0.0055 (-0.0011, 0.014) |
|  | CM | Old | 0.0012 (0.00024, 0.0079) | 0.0011 (0.00022, 0.0061) | -0.000053 (-0.0038, 0.006) |
|  | EM | Young | 0.011 (0.0012, 0.023) | 0.0073 (0.0007, 0.01) | 0.0075 (-0.0057, 0.018) |
|  | EM | Old | 0.011 (0.0022, 0.014) | 0.0072 (0.00064, 0.01) | 0.0041 (-0.0044, 0.011) |
| Ratio of fast:slow loss rates ($\lambda_A/\lambda_B$) | CM | Young | 3.3 (0.68, 27) | 8.7 (1.8, 110) | -5.2 (-100, 17) |
|  | CM | Old | 8.9 (1.6, 92) | 17 (1.7, 96) | -1.7 (-85, 82) |
|  | EM | Young | 1.8 (0.15, 14) | 3 (0.096, 24) | -0.51 (-20, 13) |
|  | EM | Old | 1.2 (0.082, 8.8) | 9.9 (0.28, 51) | -6.1 (-46, 6.1) |
| Proportion Ki67<sup>high</sup> | CM | Young | 0.5 (0.48, 0.52) | 0.31 (0.3, 0.33) | 0.19 (0.17, 0.22) |
|  | CM | Old | 0.37 (0.36, 0.39) | 0.33 (0.32, 0.35) | 0.038 (0.014, 0.061) |
|  | EM | Young | 0.27 (0.26, 0.29) | 0.16 (0.15, 0.17) | 0.11 (0.095, 0.13) |
|  | EM | Old | 0.23 (0.22, 0.25) | 0.17 (0.16, 0.18) | 0.071 (0.054, 0.084) |
| Clonal half-life of slow (days) | CM | Young | 160 (45, 1100) | 510 (160, 5700) | -340 (-5500, 530) |
|  | CM | Old | 300 (88, 2900) | 470 (110, 3200) | -72 (-2900, 2400) |
|  | EM | Young | 94 (30, 570) | 190 (68, 990) | -96 (-940, 470) |
|  | EM | Old | 88 (49, 320) | 210 (66, 1100) | -120 (-920, 210) |

### Supporting Figures

**Figure S1.**
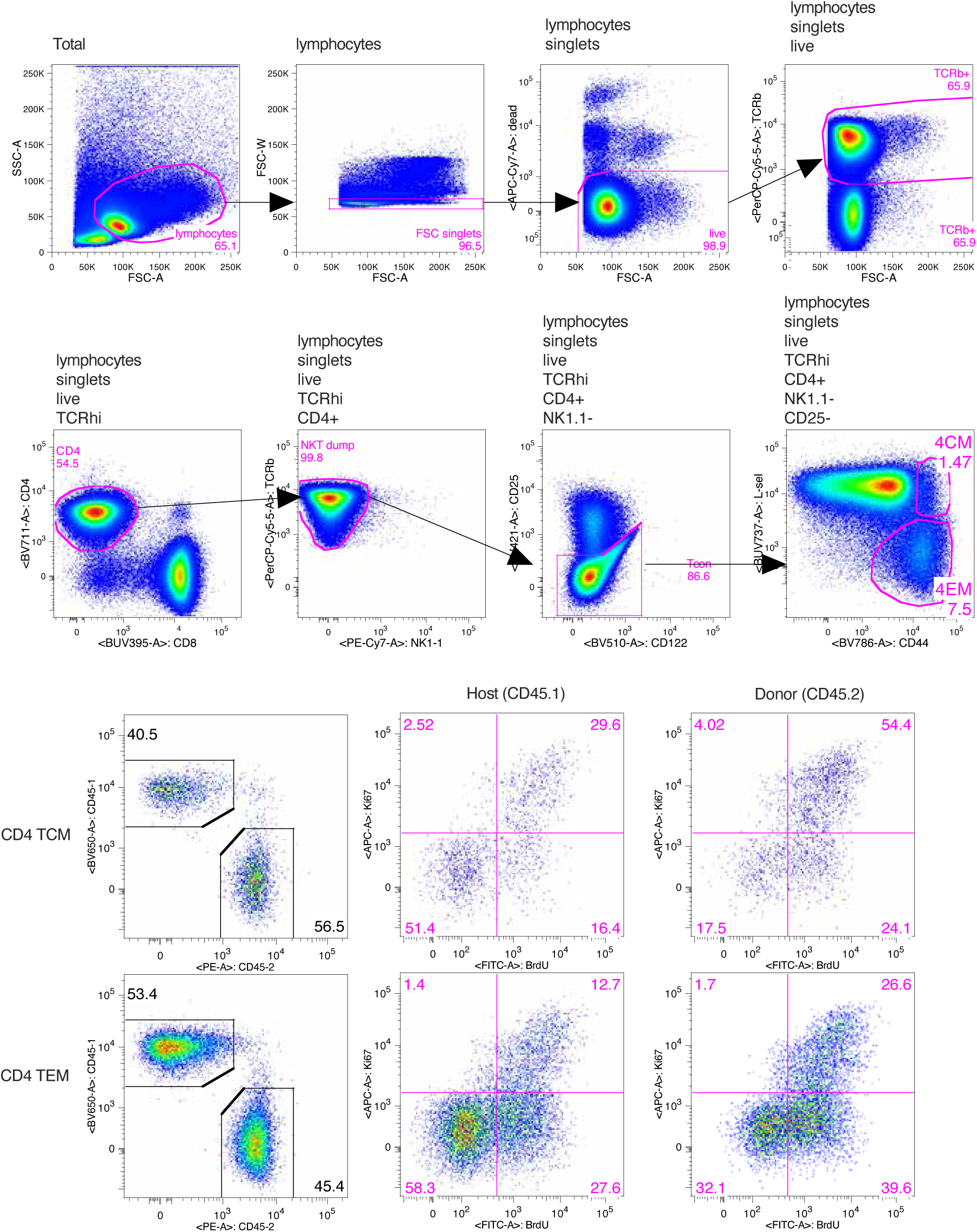
Complete gating strategy for identifying host and donor CD4 T_CM_ and T_EM_.

**Figure S2.**
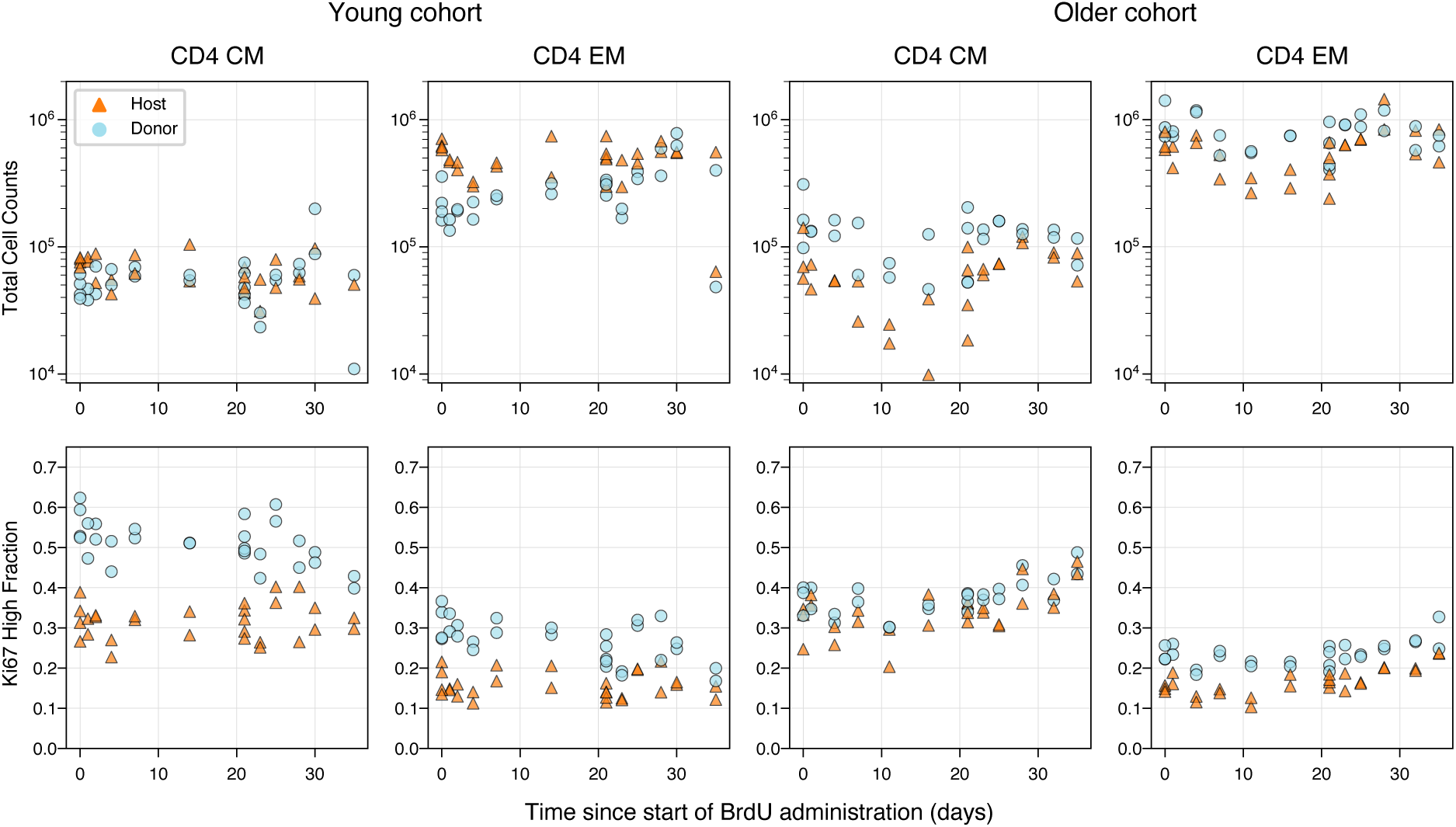
Numbers and Ki67 expression of both host and donor CD4 T_CM_ and T_EM_ over the timecourses of the labelling experiments in young and older cohorts of busulfan chimeras. The data underlying the graphs shown in the figure can be found in S1 Data.

**Figure S3.**
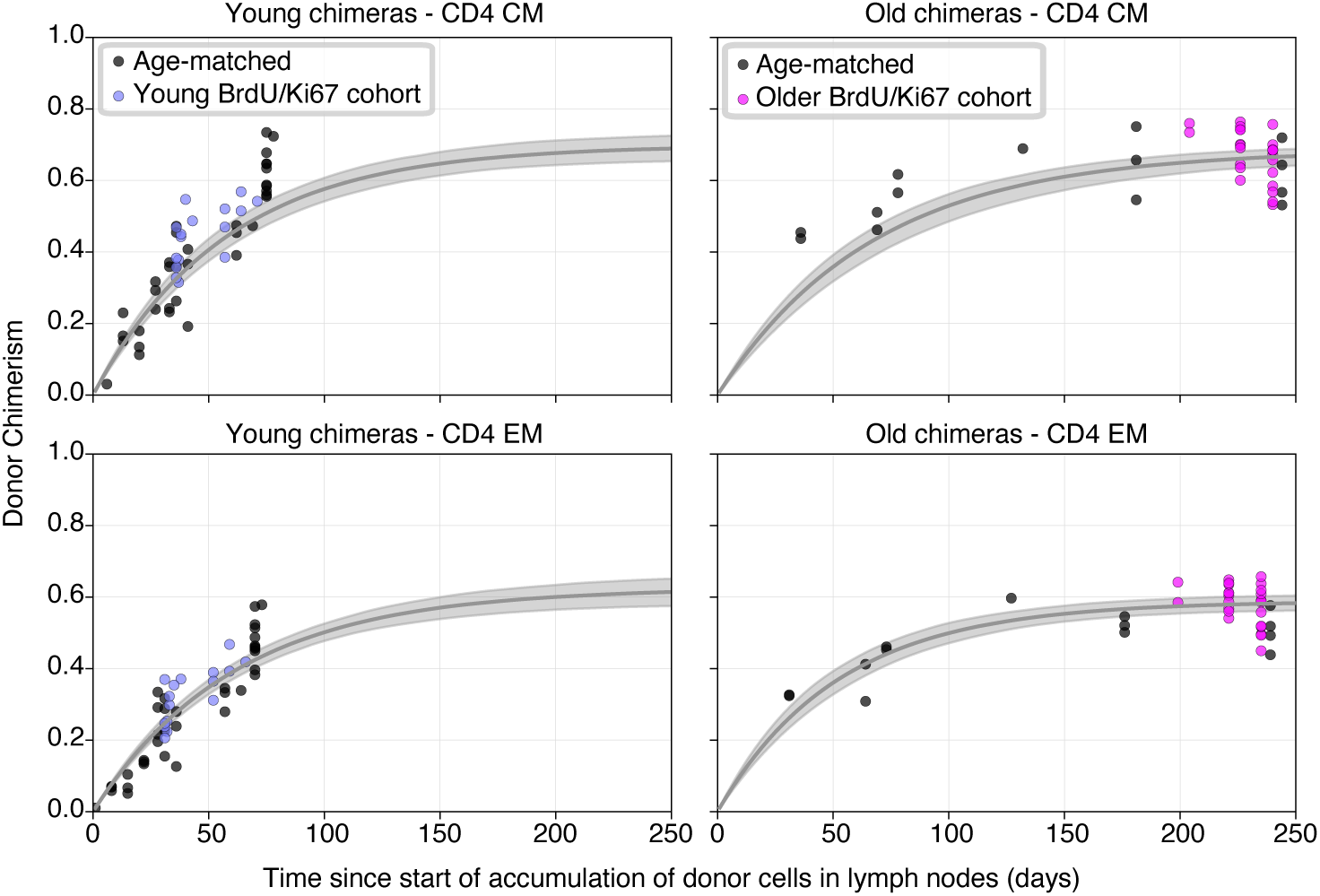
Estimating the influx into the CD4 T_CM_ and T_EM_ pools in young and old mice. Curves are the fitted trajectories of *D*(*t*)/*N*, from the solution to Eq 1 in Text C. Black points represent data from age-matched busulfan chimeric mice from other experiments. Envelopes are derived from sampling the 95% credible intervals of the parameters, and indicate where 95% of the resulting trajectories lie. The data underlying the graphs shown in the figure can be found in S1 Data.

**Figure S4.**
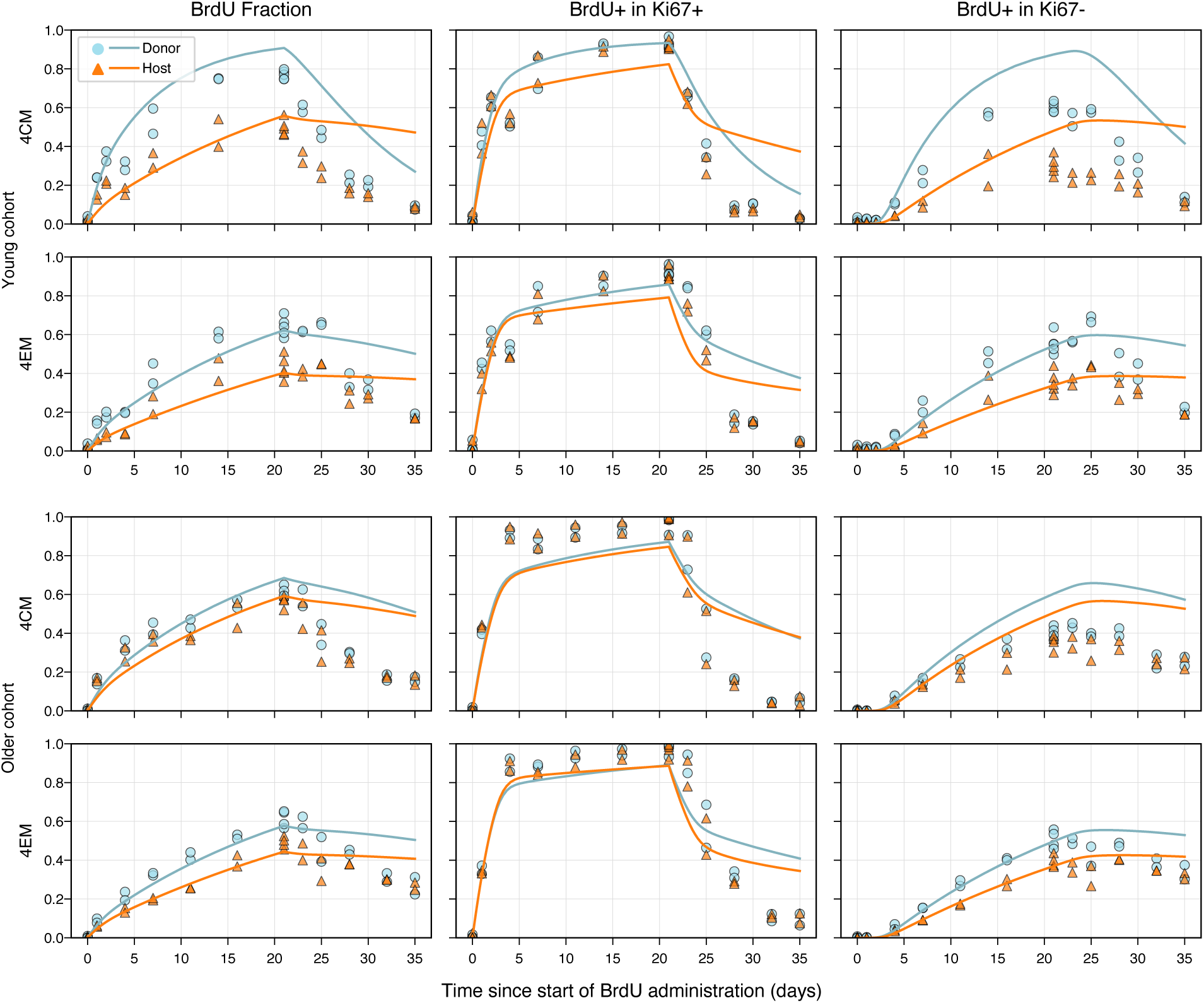
Fits of the temporal heterogeneity model to the BrdU/Ki67 labelling data. The data underlying the graphs shown in the figure can be found in S1 Data.

**Figure S5.**
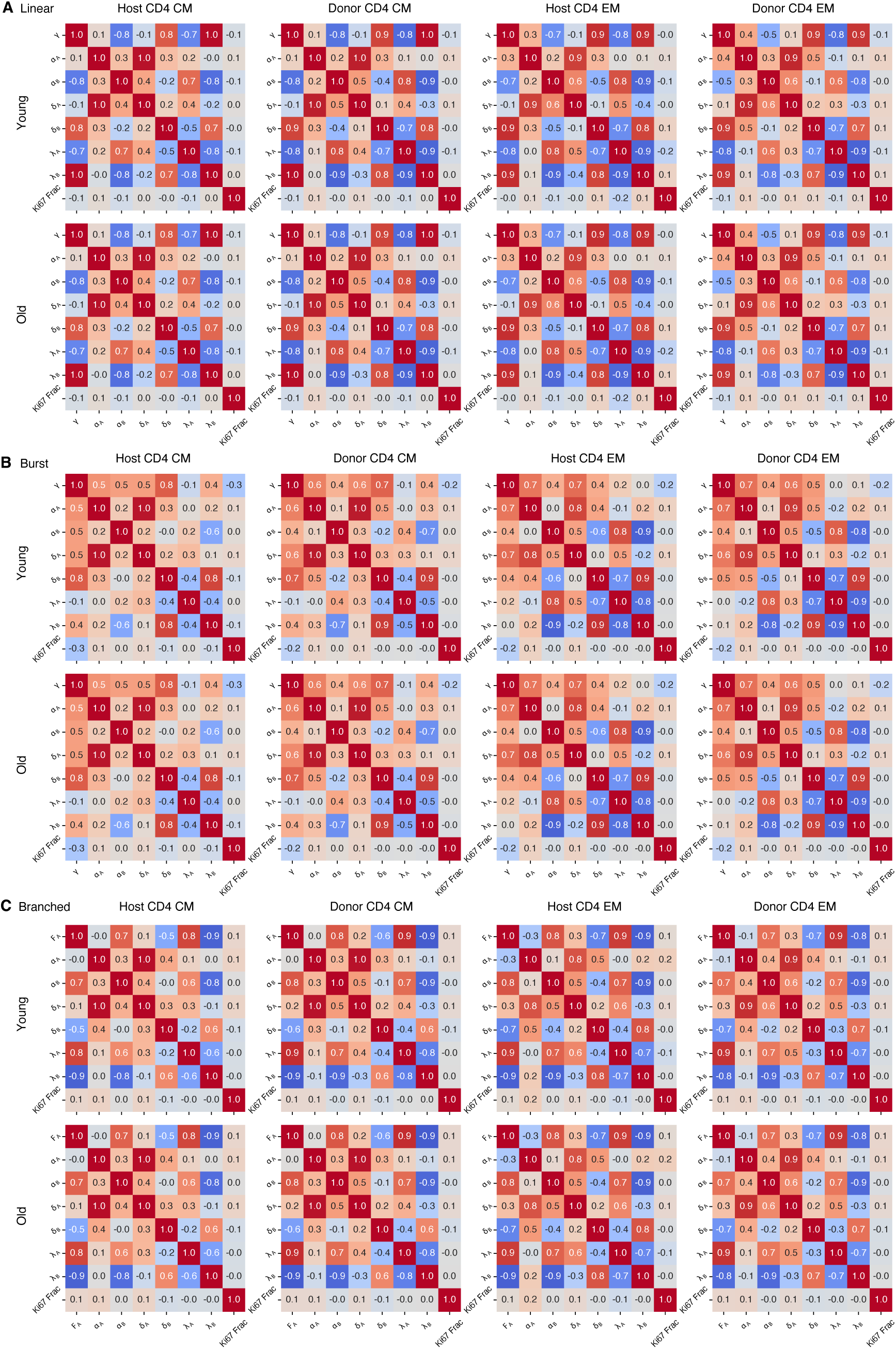
Correlations in parameters in the fits of the linear model (A), the branched model (B) and the burst model (C).

**Figure S6.**
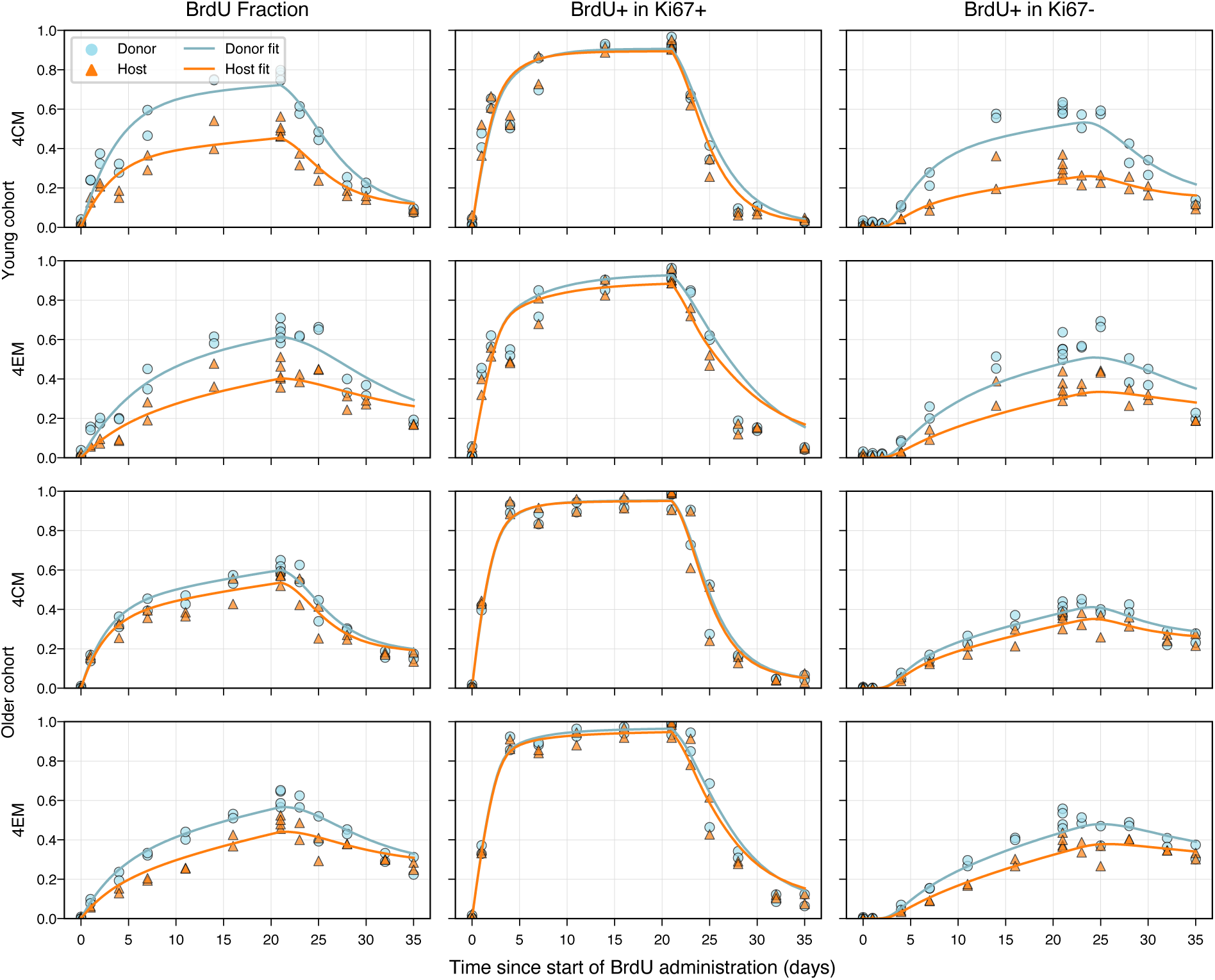
Fits of the linear model to the BrdU/Ki67 labelling data. The data underlying the graphs shown in the figure can be found in S1 Data.

**Figure S7.**
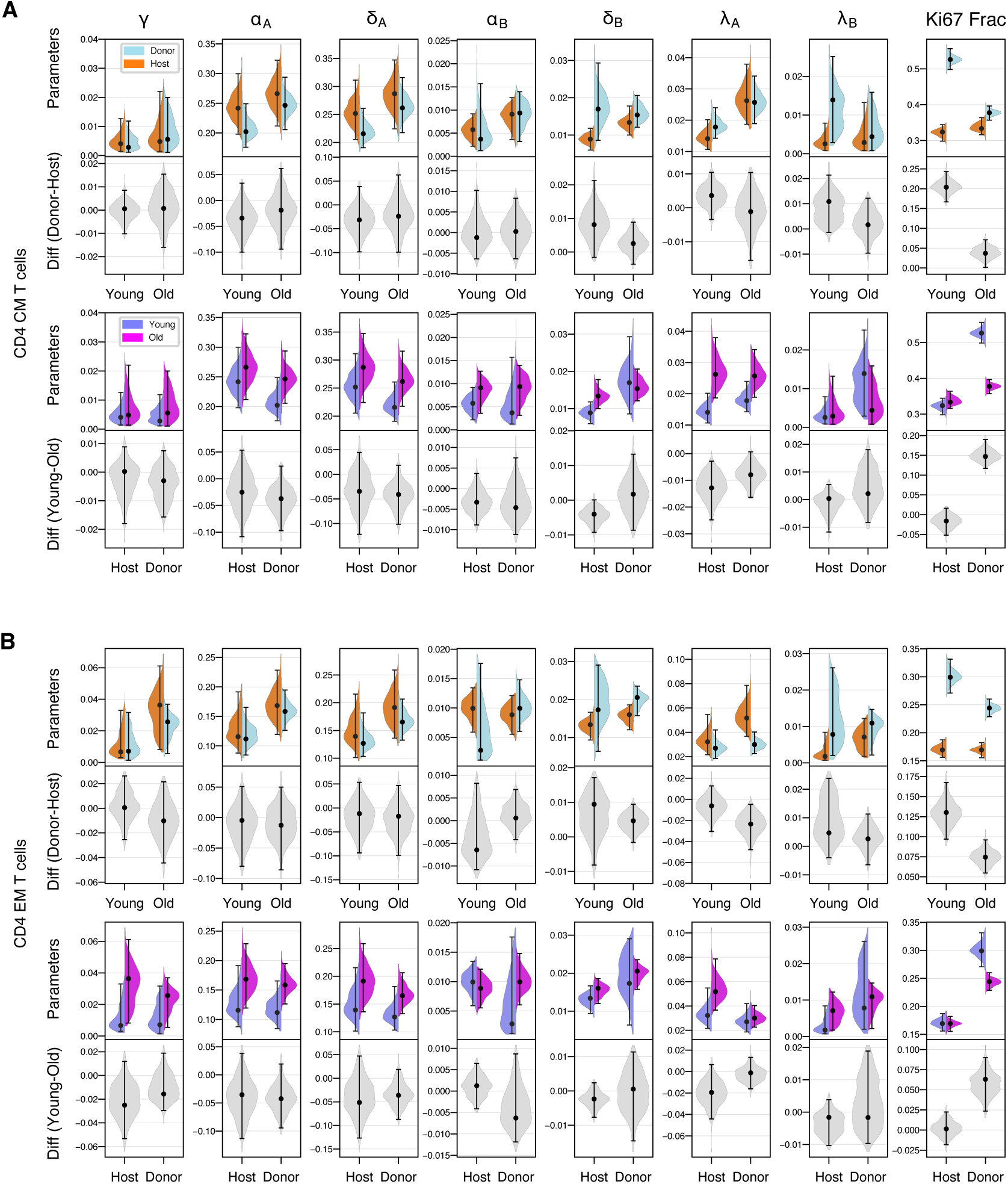
Parameter estimates for the linear model, for CD4 T_CM_ (**A**) and CD4 T_EM_ (**B**). The data underlying the graphs shown in the figure can be found in S1 Data.

**Figure S8.**
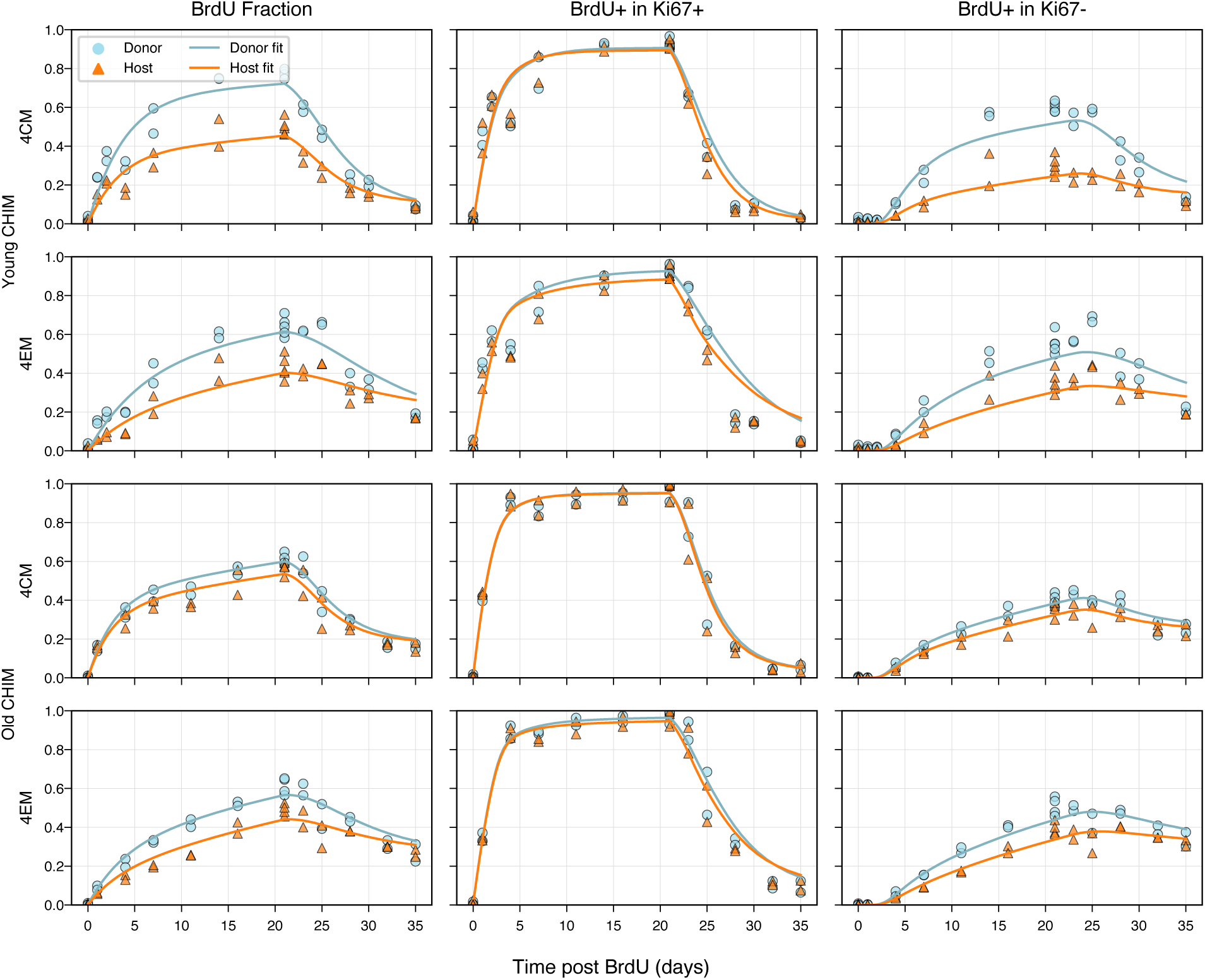
Fits of the burst model to the BrdU/Ki67 labelling data. The data underlying the graphs shown in the figure can be found in S1 Data.

**Figure S9.**
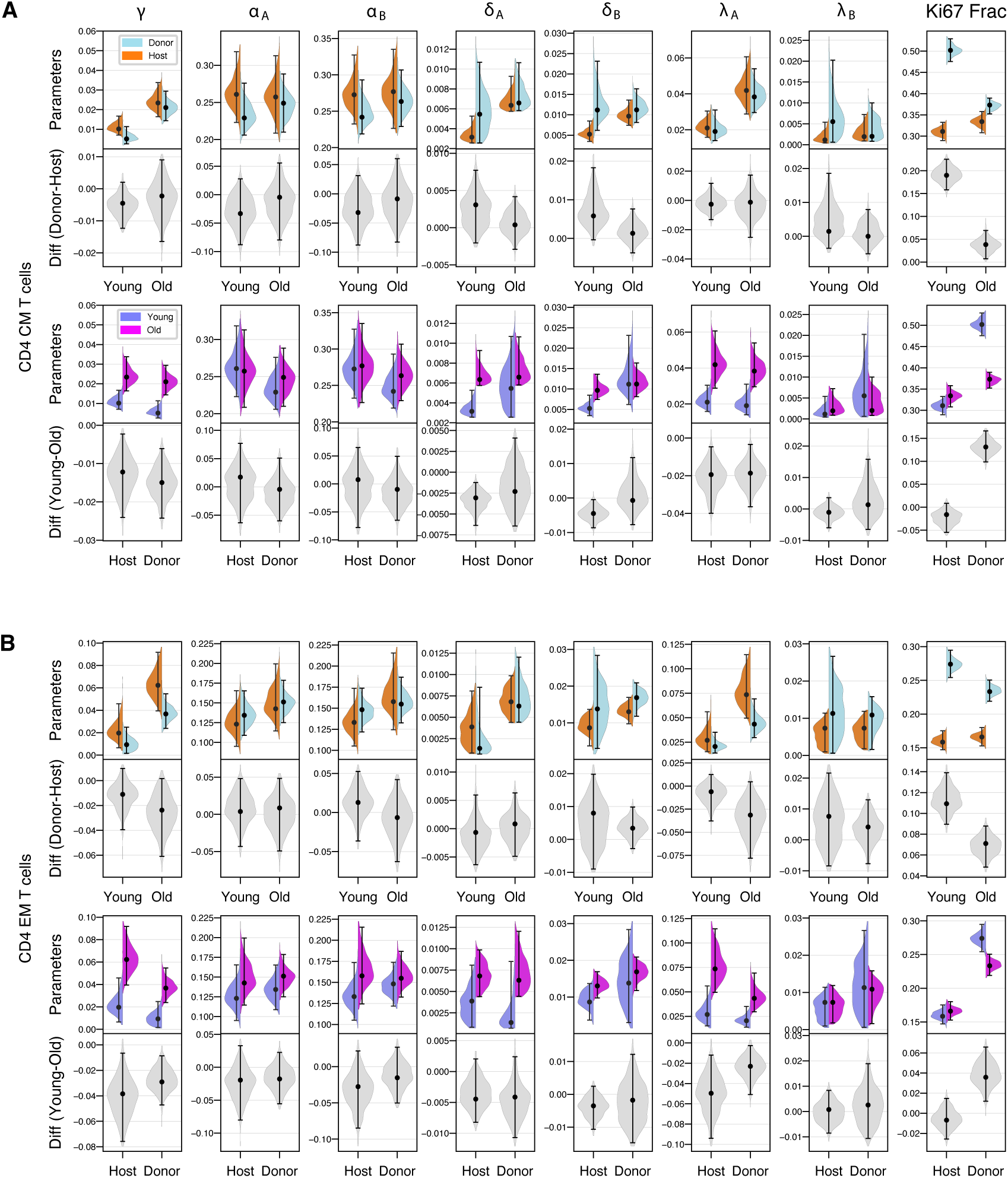
Parameter estimates for the burst model, for CD4 T_CM_ (**A**) and CD4 T_EM_ (**B**). The data underlying the graphs shown in the figure can be found in S1 Data.

**Figure S10.**
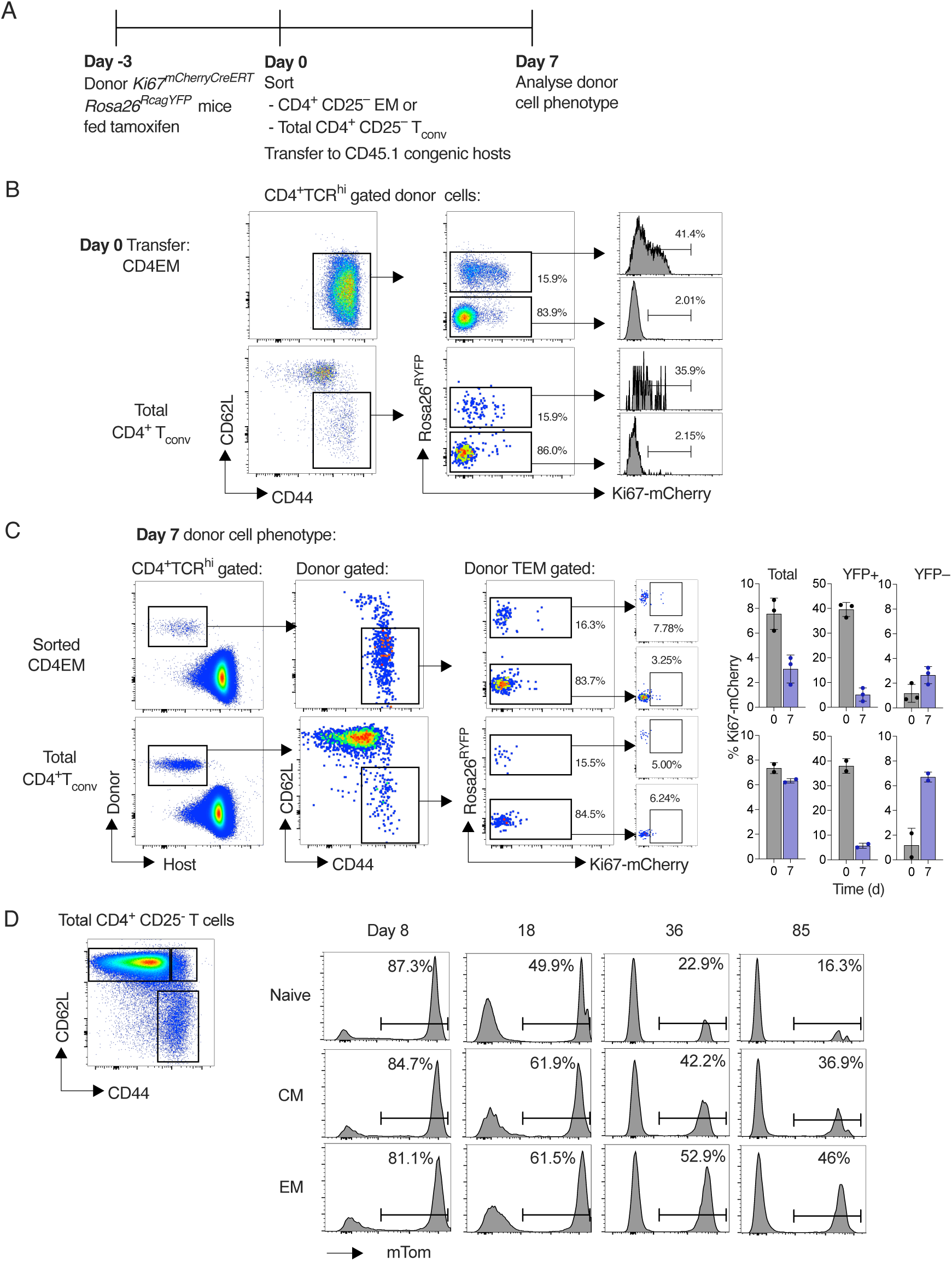
Fate mapping Ki67-expressing and CD4-expressing cells using reporter mice. (A) Ki67^mCherry-CreERT^ Rosa26^RcagYFP^ donors were fed one dose of tamoxifen (day -3). Three days later, CD4 T_EM_ or conventional CD4 T cells in bulk were purified and transferred to CD45.1 congenic hosts (day 0). At day 7, LN were recovered from hosts and donor cell phenotype analysed by flow cytometry. (B) Density plots show naive (CD44^lo^ CD62^hi^ vs. T_EM_ composition of purified populations prior to transfer, and Ki67-mcherry expression by YFP^+^ and YFP*^−^* in both donor populations prior to transfer. (C) Analysis of CD45.2^+^ CD4 T_EM_ donor populations for expression of YFP and Ki67-mCherry. Data are from three independent replicates. (D) CD4^CreERT^ Rosa26^RmTom^ reporter mice were injected with tamoxifen and culled at different days following treatment. Plots show gating strategy to identify CD4^+^ naive, T_CM_ and T_EM_ populations and examples of mTom reporter expression by these subsets at different times after tamoxifen injection. The data underlying the graphs shown in the figure can be found in S1 Data.

### Text A Mathematical modelling

#### Modelling BrdU/Ki67 pulse-chase labelling

We used ordinary differential equation models to describe the fluxes of cells between the BrdU^+/^*^−^* Ki67^high/low^ populations within the CD4 T_CM_ and T_EM_ subsets. Prior distributions on the rates of flow of new cells into each subset (*θ*) were chosen to be the posterior distributions on *θ* obtained from modelling the accumulation of donor cells in busulfan chimeric mice, as described in Text C. We assumed that kinetic heterogeneity within each subset could be represented with two subpopulations with distinct rates of division and death. In the branched model, these subpopulations (denoted A and B) are fed separately from the same precursor, at total rate *θ*, in unknown proportions. The linear model assumes constant flow from source into population A, which then transitions into population B at per capita rate *γ*. The burst model also assumes that the precursors enter population A, and cells transition into population B at per capita rate *γ*, and re-enter population A upon division. In all models, cells from from the precursor population are assumed to be Ki67^high^, based on the assumption that new memory T cells have recently undergone clonal expansion.

Levels of Ki67 protein decline continuously after mitosis but, as is standard in flow cytometry analysis, cells are categorized into binary states Ki67^high^ and Ki67^low^. We therefore modelled the transition between these two states using intermediate Ki67^high^ compartments, such that the residence time within the Ki67^high^ compartment is gamma-distributed. During the labelling phase, the delay before BrdU^+^Ki67^low^ cells appeared was clearly apparent, suggesting that the variance in the residence time within Ki67^high^ was small. To describe this kinetic we used 10 intermediate Ki67^high^ states. We also used two intermediate BrdU compartments, reflecting the assumption that in the absence of label, BrdU^+^ cells become BrdU*^−^* after two divisions. We also assumed that in the delabelling phase the BrdU content of the source population declined exponentially at a rate *µ*, which was also estimated.

The following equations describe the BrdU labelling and de-labelling phases for linear and branched models. For the branched model, the parameter *F_A_* denotes the fraction of the influx that enters subset A, while *γ* (the rate at which type A cells differentiate into type B) was set to zero. For the linear and burst models, *γ* was a free parameter, while *F_A_* was set to one. The only structural difference between the linear and burst models is that in the burst model, division of type B results in a transition back to type A.

##### Labelling phase

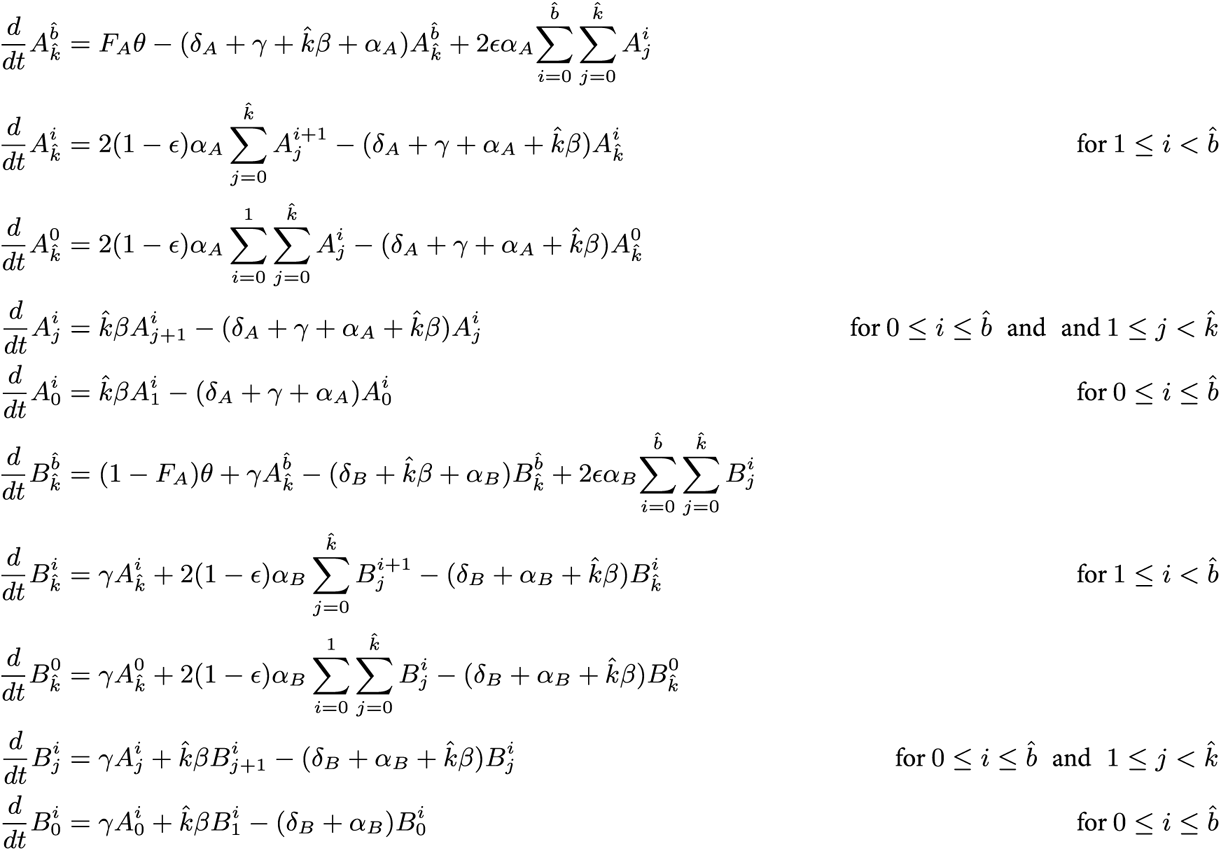

##### De-labelling phase

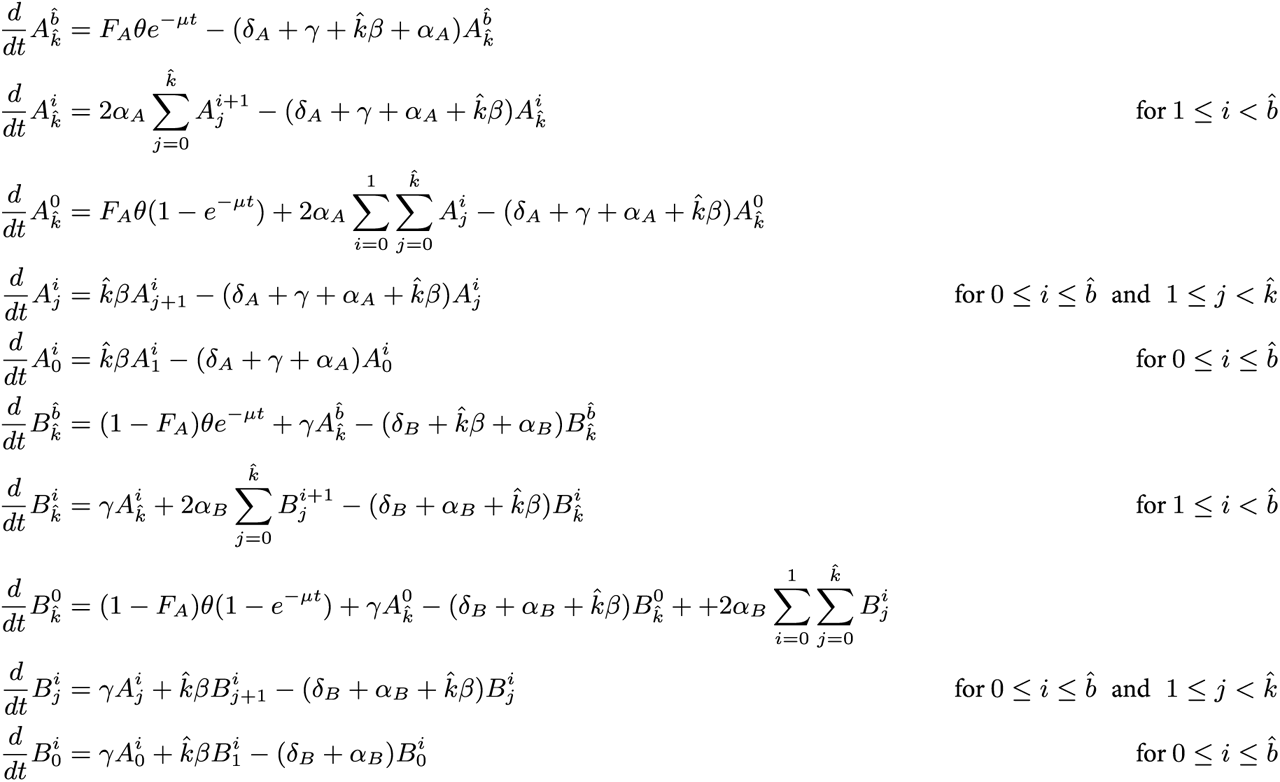

### Text B Model fitting

#### Initialisation

Motivated by the experimental observations that cells numbers and Ki67 expression levels were approximately constant over the labelling assay, we assumed that each memory population (T_CM_ and T_EM_, host/donor) was in (quasi-) equilibrium. To establish initial conditions for the model for each sampled set of parameters, we set all BrdU-labelled species to zero, and used the rk45 ODE solver to 10000 days to allow the system to attain steady state. This initialisation step determined the initial sizes of all species prior to BrdU administration, for each set of model parameters.

#### Modelling noise/mouse-to-mouse variation

The model yields total cell numbers and the proportions of cells in the four quadrants of BrdU*^±^* Ki67*^±^*. We assumed that uncertainty in the total numbers was lognormally distributed, and used a Dirichlet multinomial model to describe the assignment of cells to each of the four quadrants. The Dirichlet multinomial distribution is a discrete multivariate distribution for *k* variables *x*_1_ *. . . x_k_* where each *x_i_ ∈* (0, 1). Uncertainty in the assignment of cells to quadrants – which principally reflects variation between mice in the frequencies of cells in each quadrant – was described with an overdispersion parameter *ϕ* with an exponential prior.

#### Definition of priors

For consistency of interpretation, we defined the subsets’ loss rates (*δ_A_* and *δ_B_*) and division rates (*α_A_* and *α_B_*) such that the rates defining subset A were greater than those defining B. These rate constants were then sampled from a simplex that ensure all populations remained positive and finite in size. In addition each loss rate *δ* was constrained to be greater than the division rate to ensure the population remained at steady state with an influx of cells. The Ki67 lifetime 1/*β* was constrained to lie within 3-4 days, based on our previous studies. We chose broad priors for the rate of loss of BrdU within the source (*µ*) in the delabelling phase, and the efficiency of BrdU uptake per cell division during the labelling phase (*ϵ*). Details of priors are given at github.com/elisebullock/tcellmemorypaper.

### Text C Estimating the rates of constitutive influx into CD4 T_CM_ and T_EM_

Within busulfan chimeric mice, the accumulation of donor-derived T_CM_ or T_EM_ (Fig S3) was reasonably well described by a simple model with a single average rate of turnover, *λ*;

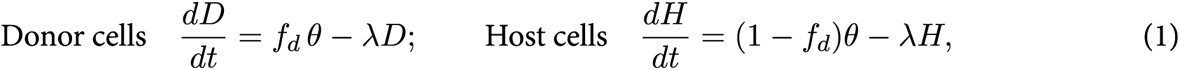

where *f_d_* is the proportion of the constitutive influx of memory *θ* that comprises donor cells, and we model from the time at which donor memory T cells start to appear in lymph nodes. For each subset (T_CM_ and T_EM_) and in young and old mice, we fitted this model simultaneously to the steady state numbers *N* = *D* + *H* = *θ*/*λ* and the chimerism timecourse *D*(*t*)/*N* (Fig S3), using data from the chimeric mice we studied here augmented with data from age-matched chimeric mice aggregated from other experiments. This procedure yielded posterior distributions of the total (host plus donor) influx rates *θ* (Table S1) that were only weakly correlated with those of the other parameters (*N*, *λ*, and *f_d_*).

Text D Explaining the progressive loss of Ki67 in donor cells with mouse age and the convergence of host and donor cell dynamics in older mice

To understand the patterns of host/donor cell differences at the level of their population dynamics, we derived an expression for the proportion of a memory population that is fast cells, *p_A_*, in the branched model:

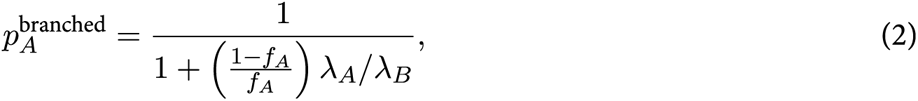

where *f_A_* is the proportion of the source entering the fast population, which was slightly lower for T_EM_ than T_CM_ (Fig 4 in the main text). For both subsets we saw no significant differences in *f_A_* between host and donor cells or between young and old mice, indicating that for each subset the rationing of new memory cells into fast and slow subsets was an invariant quantity. The differences in the donor/host fast fractions must therefore derive from the ratio of the loss rates of fast to slow populations, *λ_A_*/*λ_B_*. In young mice, fast donor T_CM_ clones were marginally less persistent than host counterparts 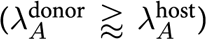 but slow donor clones were much less persistent than slow host clones 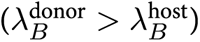. Thus, for T_CM_, *λ_A_*/*λ_B_* was smaller for donor than host, leading to a greater fast cell fraction and so explaining the donor-skewed Ki67 fraction (Fig 2D in the main text). For T_EM_ the picture was similar; in the younger cohort, clonal lifespans of fast cells were similar for donor and host, but slow donor clones were substantially shorter lived than host clones. Therefore, again, *λ_A_*/*λ_B_* was smaller for donor than host. In older mice, all differences in parameters between host and donor cells shrank, although the fast fraction remained significantly higher for donor in both T_CM_ and T_EM_. These patterns reflected the degree of convergence of Ki67 expression of host and donor cells (Fig 2D).

These conclusions also held for the linear and burst models. For both, the fraction of the population made up of fast cells is given by the simpler expression

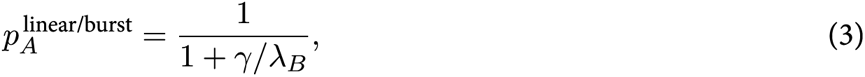

where *γ* is the rate of differentiation from fast (A) to slow (B). For the linear model, for both T_CM_ and T_EM_, we saw no differences in *γ* between host and donor, or across mouse age (Fig S7). In contrast, *λ_B_*, the loss rate of slow clones, was greater for donor than for host in the younger mice, again explaining the higher fast fraction among donor cells in young mice, and hence their higher Ki67 expression. In older mice, *λ_B_* for donor cells approached that of host, yielding the same conclusion drawn from the branched model. For the burst model, the same trends held for *λ_B_* (Fig S9). There was an additional contribution from *γ*, the rate of return of fast cells to a more quiescent state. In younger mice, this rate was estimated to be slightly lower for donor than for host cells (that is, younger memory cells ‘burst’ for slightly longer on average than older cells); this difference also acted to increase the fast fraction *p_A_* among donor cells.

### Text E Calculation of mean lifespan in the branched model

Suppose the donor fraction is *f_D_*, the fast cell fractions within host and donor are 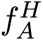 and 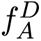, respectively, and 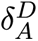 and 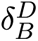 are the loss rates of fast and slow donor cells, respectively, with correspondingly 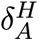 and 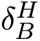 for host cells. Then the population-average loss rate 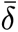 is given by

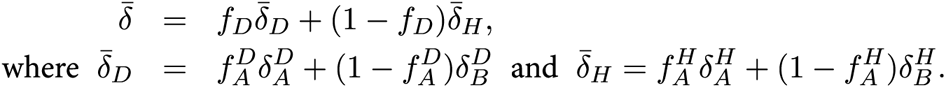

The mean lifespan is then 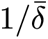.

### Text F Obtaining mean lifespans through estimating the total production rate

We present two methods of obtaining an approximate average loss rate (and hence mean lifespan) by measuring the total rate of production. These are equal for a population at equilibrium.

The first is to use information contained in the initial growth in the BrdU labelled fraction. De Boer and Perelson [1] used data from Younes *et al.* [2] who found that 35% and 60% of memory CD4 T cells were BrdU^+^ after 3 and 10 days of labelling, respectively, to estimate an expected life span of between 14 and 22 days. To generalise their modelling approach, consider a memory population of size *N* at equilibrium, fed from a source at rate *θ*, dividing at average rate *α*, and lost at rate *δ*. During BrdU administration, assume the source is entirely labelled, and unlabelled memory cells take up BrdU with probability *ϵ* per division. If we denote unlabelled cells *N_U_* and labelled cells *N_L_*, then

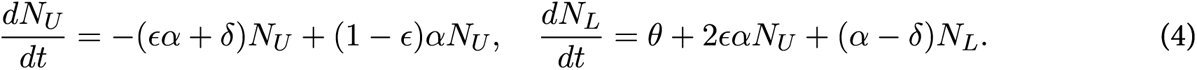

The labelled fraction *L* = *N_L_*/*N* is given by

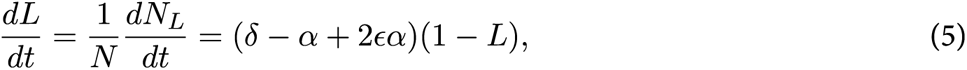

where we have used *N_U_* = *N − N_L_*, and the steady state condition *dN* /*dt* = 0 yielding *θ*/*N* = *δ − α*. The initial upslope of the labelled fraction *L* is then *p* = *δ − α* + 2*ϵα*. If the source is entirely unlabelled,

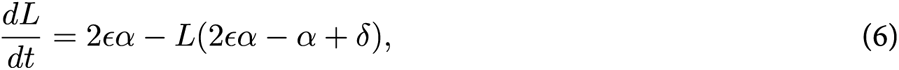

and so *p* = 2*ϵα*. Therefore, depending on the rate of influx and its label content, the initial upslope in the BrdU^+^ fraction can be a combination of both the average division and loss rates, as well as the uptake efficiency.

De Boer and Perelson’s calculation assumed that memory CD4 T cells are entirely self-renewing, such that *θ* = 0 and *α* = *δ*, and that uptake of BrdU by dividing cells is 100% efficient. Eq 6 then yields *dL*/*dt* = 2*δ*(1*−L*), which they fitted to the Younes data to obtain the average lifespan 1/*δ*. The self-renewing approximation is reasonable – we estimate that the influxes into T_CM_ and T_EM_ are of the order 1% of the population size per day, so we can proceed by neglecting them. However we established that the BrdU uptake efficiency *ϵ* is considerably less than 1, and was well constrained by the steepness of growth in the BrdU^+^Ki67^high^ fraction (*ϵ* = 0.55 in younger cohort, and 0.75 in the older cohort, which were separate experiments). The mean lifespan is then reduced to 2*ϵ*/*p*, as quoted in the main text.

The second approach is to derive the average rate of turnover using the frequency of cells expressing Ki67. Consider again the memory population at steady state size *N* with cells entering at rate *θ*, dividing at an average rate *α* and dying at average rate *δ*:

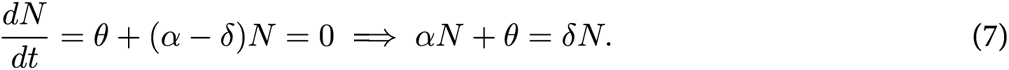

We know that Ki67 is expressed for a time *T ≃* 3 days, and we assume that all cells entering memory are Ki67^high^. Then the number of cells in the population at any time *t* that are Ki67^high^, *K*^+^, is equal to the number that entered or divided in the last *T* days, and survived to time *t*:

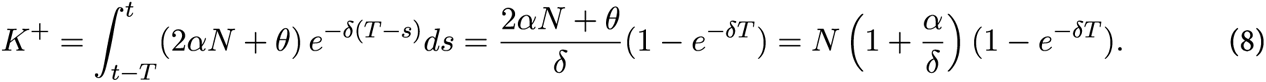

If the influx per unit time makes up a small proportion of the population (*θ ≪ N*), as we found for both T_CM_ and T_EM_, then *θ ≃* 0 and *α ≃ δ*. Then using Eq 7, the Ki67^high^ proportion *k* = *K*^+^/*N ≃* 2(1 *− e^−δT^*). The mean lifespan *τ* = 1/*δ* is then

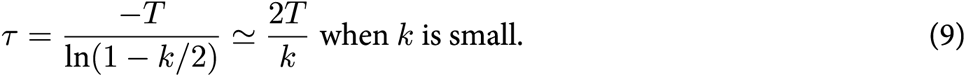

### Text G Predicting patterns of Ki67 expression among YFP^+^ and YFP*^−^* cells following adoptive transfer

**Figure A.**
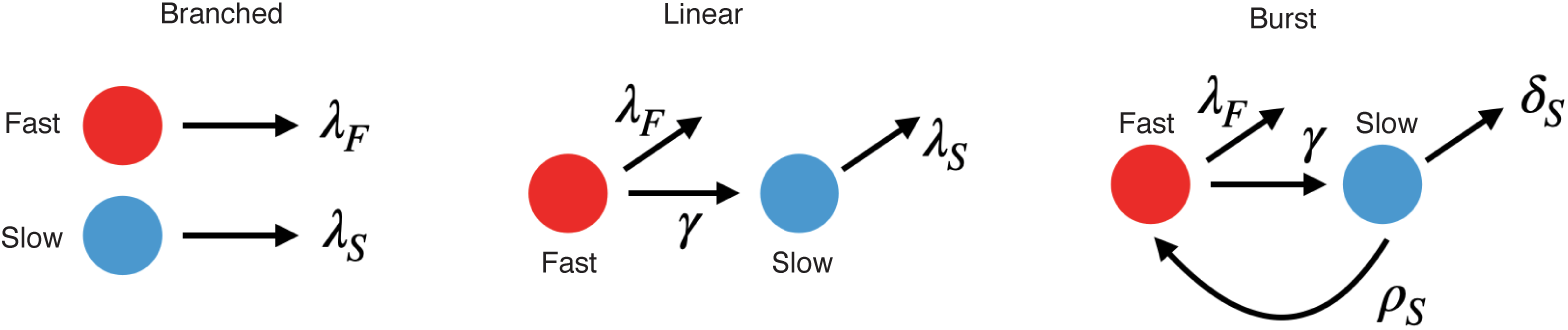
Schematic of the three models, indicating the key parameters.

We can use each model to predict the outcome of transferring fast and slow cells to a congenic host after labelling Ki67-expressing cells with YFP. First, we note that in all models, we estimate that fast clones are lost more rapidly than slow (*λ_F_ > λ_S_*).

The branched model predicts that slow cells, which are largely YFP*^−^* but will nevertheless contain some YFP^+^ cells, will come to dominate over time. Therefore, Ki67 in both YFP^+^ and YFP*^−^* populations will fall, and converge.

In the linear model, the slow cells will comprise cells that were slow upon transfer, and cells that subsequently transitioned from fast. Since *λ_F_ > λ_S_*, selection will occur within the slow population for cells that were originally slow. So again, Ki67 overall falls to a lower level, that is independent of YFP expression.

Finally, in the burst model, there is a period just after transfer when YFP^+^ cells are almost all transiently fast, and Ki67^high^. Without new fast cells flowing in, the system will then settle to a lower level of Ki67 expression. YFP^+^ and YFP*^−^* cells then behave identically, because all have the potential to re-enter the burst phase at random.

In summary, all three models predict the outcome of the transfer experiment.

### Text H Predicting the timecourses of mTomato expression in CD4 reporter mice

We generated predicted timecourses of mTom-expressing cell frequencies as follows. First, we assumed that naive CD4 T cells were the dominant precursor of T_CM_. We then fitted a phenomenological power-law function to the timecourse of mTom^+^ cells starting at day 8 after tamoxifen treatment, at which point mTom expression was maximal in all compartments (Fig 7C in the main text). This function defined the time-dependent source population for mTom^+^ CD4 T_CM_. We assumed that mTom^+^ and mTom*^−^* cells transitioned from CD4 naive to T_CM_ at the same rates. Second, we initialized the populations of Ki67-high, intermediate and low CD4 T_CM_ and T_EM_ within mTom^+^ and mTom*^−^* cells, using the observed overall Ki67 fraction; and distributed these between fast and slow cells using the fast fraction derived from our fitted parameters (Text D). This process yielded the initial numbers of cells within all subpopulations (Text A). Then, using the parameters derived from fitting the BrdU/Ki67 timecourses, and the function describing the mTom^+^ fraction among naive CD4 T cells, we simulated 150-day time courses of the mTom^+^ fractions within T_CM_ and T_EM_ (Fig 7D in the main text), considering both naive or T_CM_ as possible sources for T_EM_.

## References

1. Swain SL, Agrewala JN, Brown DM, Jelley-Gibbs DM, Golech S, Huston G, et al. CD4+ T-cell memory: generation and multi-faceted roles for CD4+ T cells in protective immunity to influenza. Immunological reviews. 2006;211(1):8–22.

2. Tubo NJ, Jenkins MK. CD4+ T Cells: guardians of the phagosome. Clin Microbiol Rev. 2014;27(2):200–13. doi:10.1128/CMR.00097-13.

3. Künzli M, Masopust D. CD4+ T cell memory. Nature Immunology. 2023;16:4. doi:10.1038/s41590-023-01510-4.

4. Westera L, Drylewicz J, den Braber I, Mugwagwa T, van der Maas I, Kwast L, et al. Closing the gap between T-cell life span estimates from stable isotope-labeling studies in mice and humans. Blood. 2013;122(13):2205–12. doi:10.1182/blood-2013-03-488411.

5. Gossel G, Hogan T, Cownden D, Seddon B, Yates AJ. Memory CD4 T cell subsets are kinetically heterogeneous and replenished from naive T cells at high levels. eLife. 2017;6:e23013.

6. Baliu-Piqué M, Verheij MW, Drylewicz J, Ravesloot L, de Boer RJ, Koets A, et al. Short Lifespans of Memory T-cells in Bone Marrow, Blood, and Lymph Nodes Suggest That T-cell Memory Is Maintained by Continuous Self-Renewal of Recirculating Cells. Front Immunol. 2018;9:2054.

7. Borghans JAM, Tesselaar K, de Boer RJ. Current best estimates for the average lifespans of mouse and human leukocytes: reviewing two decades of deuterium-labeling experiments. Immunol Rev. 2018;285(1):233–248.

8. Baliu-Piqué M, Otto SA, Borghans JAM, Tesselaar K. In vivo deuterium labelling in mice supports a dynamic model for memory T-cell maintenance in the bone marrow. Immunol Lett. 2019;210:29–32.

9. Derksen LY, Tesselaar K, Borghans JAM. Memories that last: Dynamics of memory T cells throughout the body. Immunol Rev. 2023;316(1):38–51. doi:10.1111/imr.13211.

10. del Amo PC, Beneytez JL, Boelen L, Ahmed R, Miners KL, Zhang Y, et al. Human TSCM cell dynamics in vivo are compatible with long-lived immunological memory and stemness. PLoS Biology. 2018;16(6):1–22.

11. Jameson SC, Lee YJ, Hogquist KA. Innate Memory T cells. In: Advances in Immunology. vol. 126. Adv Immunol; 2015. p. 173–213.

12. Kawabe T, Jankovic D, Kawabe S, Huang Y, Lee PHH, Yamane H, et al. Memory-phenotype CD4+ T cells spontaneously generated under steady state conditions exert innate Th1-like effector function. Science Immunology. 2017;2(12):16.

13. Min B, Paul WE. Endogenous proliferation: burst-like CD4 T cell proliferation in lymphopenic settings. Semin Immunol. 2005;17(3):201–7. doi:10.1016/j.smim.2005.02.005.

14. Tough DF, Sprent J. Turnover of naive- and memory-phenotype T cells. J Exp Med. 1994;179(4):1127–35.

15. Lenz DC, Kurz SK, Lemmens E, Schoenberger SP, Sprent J, Oldstone MB, et al. IL-7 regulates basal homeostatic proliferation of antiviral CD4+T cell memory. Proc Natl Acad Sci U S A. 2004;101(25):9357–62.

16. Purton JF, Tan JT, Rubinstein MP, Kim DM, Sprent J, Surh CD. Antiviral CD4+ memory T cells are IL-15 dependent. J Exp Med. 2007;204(4):951–61.

17. Younes SA, Punkosdy G, Caucheteux S, Chen T, Grossman Z, Paul WE. Memory phenotype CD4 T cells undergoing rapid, nonburst-like, cytokine-driven proliferation can be distinguished from antigen-experienced memory cells. PLoS Biology. 2011;9(10):e1001171.

18. Hogan T, Nowicka M, Cownden D, Pearson CF, Yates AJ, Seddon B. Differential impact of self and environmental antigens on the ontogeny and maintenance of CD4+ T cell memory. eLife. 2019;8. doi:10.7554/eLife.48901.

19. Mohri H, Perelson AS, Tung K, Ribeiro RM, Ramratnam B, Markowitz M, et al. Increased turnover of T lymphocytes in HIV-1 infection and its reduction by antiretroviral therapy. J Exp Med. 2001;194(9):1277–87.

20. Asquith B, Debacq C, Macallan DC, Willems L, Bangham CRM. Lymphocyte kinetics: the interpretation of labelling data. Trends in Immunology. 2002;23(12):596–601.

21. Ganusov VV, Borghans JAM, De Boer RJ. Explicit kinetic heterogeneity: mathematical models for interpretation of deuterium labeling of heterogeneous cell populations. PLoS Comput Biol. 2010;6(2):e1000666.

22. De Boer RJ, Perelson AS, Ribeiro RM. Modelling deuterium labelling of lymphocytes with temporal and/or kinetic heterogeneity. Journal of the Royal Society Interface. 2012;9(74):2191–2200.

23. Hogan T, Gossel G, Yates AJ, Seddon B. Temporal fate mapping reveals age-linked heterogeneity in naive T lymphocytes in mice. Proceedings of the National Academy of Sciences. 2015;112(50):E6917—-E6926. doi:10.1073/pnas.1517246112.

24. Hogan T, Yates A, Seddon B. Generation of Busulfan chimeric mice for the analysis of T cell population dynamics. Bio-protocol. 2017;4(24).

25. Hogan T, Yates A, Seddon B. Analysing temporal dynamics of T cell division *in vivo* using Ki67 and BrdU co-labelling by flow cytometry. Bio-protocol. 2017;7(24).

26. Verheijen M, Rane S, Pearson C, Yates AJ, Seddon B. Fate Mapping Quantifies the Dynamics of B Cell Development and Activation throughout Life. Cell Reports. 2020;33(7):108376. doi:10.1016/j.celrep.2020.108376.

27. Rane S, Hogan T, Seddon B, Yates AJ. Age is not just a number: Naive T cells increase their ability to persist in the circulation over time. PLoS biology. 2018;16(4):e2003949. doi:10.1371/journal.pbio.2003949.

28. Rane S, Hogan T, Lee E, Seddon B, Yates AJ. Towards a unified model of naive T cell dynamics across the lifespan. eLife. 2022;11:11. doi:10.7554/eLife.78168.

29. Lukas E, Hogan T, Williams C, Seddon B, Yates AJ. Quantifying cellular dynamics in mice using a novel fluorescent division reporter system. Frontiers in Immunology. 2023;14. doi:10.3389/fimmu.2023.1157705.

30. Sasaki K, Murakami T, Kawasaki M, Takahashi M. The cell cycle associated change of the Ki-67 reactive nuclear antigen expression. Journal of Cellular Physiology. 1987;133(3):579–584.

31. Miller I, Min M, Yang C, Tian C, Gookin S, Carter D, et al. Ki67 is a graded rather than a binary marker of proliferation versus quiescence. Cell Rep. 2018;24(5):1105–1112.e5.

32. De Boer RJ, Yates AJ. Modeling T Cell Fate. Annu Rev Immunol. 2023;41:513–532. doi:10.1146/annurev-immunol-101721-040924.

33. Costa Del Amo P, Debebe B, Razavi-Mohseni M, Nakaoka S, Worth A, Wallace D, et al. The Rules of Human T Cell Fate in vivo. Front Immunol. 2020;11:573. doi:10.3389/fimmu.2020.00573.

34. De Boer RJ, Perelson AS. Quantifying T lymphocyte turnover. Journal of Theoretical Biology. 2013;327:45–87.

35. Schittler D, Allgöwer F, De Boer RJ. A new model to simulate and analyze proliferating cell populations in BrdU labeling experiments. BMC Syst Biol. 2013;7 Suppl 1:S4.

36. Tsukamoto H, Clise-Dwyer K, Huston GE, Duso DK, Buck AL, Johnson LL, et al. Age-associated increase in lifespan of naive CD4 T cells contributes to T-cell homeostasis but facilitates development of functional defects. Proc Natl Acad Sci U S A. 2009;106(43):18333–8.

37. Mold JE, Réu P, Olin A, Bernard S, Michaëlsson J, Rane S, et al. Cell generation dynamics underlying naive T-cell homeostasis in adult humans. PLoS biology. 2019;17(10). doi:10.1371/JOURNAL.PBIO.3000383.

38. Swain AC, Borghans JAM, de Boer RJ. Effect of cellular aging on memory T-cell homeostasis. Front Immunol. 2022;13:947242. doi:10.3389/fimmu.2022.947242.

39. Linton PJ, Haynes L, Tsui L, Zhang X, Swain S. From naive to effector–alterations with aging. Immunol Rev. 1997;160:9–18. doi:10.1111/j.1600-065x.1997.tb01023.x.

40. Haynes L, Linton PJ, Eaton SM, Tonkonogy SL, Swain SL. Interleukin 2, but not other common gamma chain-binding cytokines, can reverse the defect in generation of CD4 effector T cells from naive T cells of aged mice. J Exp Med. 1999;190(7):1013–24. doi:10.1084/jem.190.7.1013.

41. Haynes L, Eaton SM, Burns EM, Rincon M, Swain SL. Inflammatory cytokines overcome age-related defects in CD4 T cell responses in vivo. J Immunol. 2004;172(9):5194–9. doi:10.4049/jimmunol.172.9.5194.

42. Akondy RS, Fitch M, Edupuganti S, Yang S, Kissick HT, Li KW, et al. Origin and differentiation of human memory CD8 T cells after vaccination. Nature. 2017;552(7685):362–367.

43. Zarnitsyna VI, Akondy RS, Ahmed H, McGuire DJ, Zarnitsyn VG, Moore M, et al. Dynamics and turnover of memory CD8 T cell responses following yellow fever vaccination. PLoS Comput Biol. 2021;17(10):e1009468. doi:10.1371/journal.pcbi.1009468.

44. Seddon B, Tomlinson P, Zamoyska R. Interleukin 7 and T cell receptor signals regulate homeostasis of CD4 memory cells. Nature Immunology. 2003;4(7):680–686.

45. Choo DK, Murali-Krishna K, Anita R, Ahmed R. Homeostatic turnover of virus-specific memory CD8 T cells occurs stochastically and is independent of CD4 T cell help. J Immunol. 2010;185(6):3436–44.

46. Pepper M, Linehan JL, Pagán AJ, Zell T, Dileepan T, Cleary PP, et al. Different routes of bacterial infection induce long-lived TH1 memory cells and short-lived TH17 cells. Nat Immunol. 2010;11(1):83–9.

47. Sledzińska A, Hemmers S, Mair F, Gorka O, Ruland J, Fairbairn L, et al. TGF-*β* signalling is required for CD4+ T cell homeostasis but dispensable for regulatory T cell function. PLoS Biol. 2013;11(10):e1001674. doi:10.1371/journal.pbio.1001674.

48. Vehtari A, Gelman A, Gabry J. Practical Bayesian model evaluation using leave-one-out cross-validation and WAIC. Statistics and Computing. 2017;27(5):1413–1432. doi:10.1007/s11222-016-9696-4.

## References

1. De Boer RJ, Perelson AS. Quantifying T lymphocyte turnover. Journal of Theoretical Biology. 2013;327:45–87.

2. Younes SA, Punkosdy G, Caucheteux S, Chen T, Grossman Z, Paul WE. Memory phenotype CD4 T cells undergoing rapid, nonburst-like, cytokine-driven proliferation can be distinguished from antigen-experienced memory cells. PLoS Biology. 2011;9(10):e1001171.

